# Somatic hypermutation-mediated paratope flexibility improves the cross-reactivity of human malaria antibodies

**DOI:** 10.1101/2024.07.24.604858

**Authors:** Rajagopal Murugan, Anton Hanke, Opeyemi Ernest Oludada, Giulia Costa, Yevel Flores-Garcia, Ghulam Mustafa, Kan Li, Richard H. C. Huntwork, Gillian Q. Horn, S. Moses Dennison, Georgia D Tomaras, Fidel Zavala, Elena A. Levashina, Randal R. Ketchem, Rebecca C. Wade, Hedda Wardemann

## Abstract

The protective capacity of antibodies targeting circumsporozoite protein on sporozoites of the malaria parasite *Plasmodium falciparum* (PfCSP) is linked to high affinity and cross-reactivity with the PfCSP central repeat domain and N-terminal junction. However, the role of somatic hypermutation (SHM) in the development of such antibodies remains unclear. Here we define the contributions of SHM to the high affinity and strong repeat and N-junction cross-reactivity of the potent anti-PfCSP monoclonal antibody (mAb) 4493 and of similar antibodies with shared SHM and affinity maturation trajectories. Molecular dynamics simulations reveal that SHM reduces the flexibility of the unbound mAb 4493 but increases the flexibility of the antigen-bound complex, thereby lowering the entropic cost for antigen binding. Furthermore, we identify an inverse relation between antibody affinity and serum stability, which limits the protective capacity of these antibodies. Our study provides molecular level evidence for the different roles that the SHM process plays in increasing VH3-49+Vκ3-20 antibody affinity and cross-reactivity and demonstrates how antibody affinity maturation can negatively impact antibody stability and thereby function.

## INTRODUCTION

*Plasmodium falciparum* (Pf) malaria remains a global health problem and a leading cause of childhood mortality, especially in Sub-Saharan Africa (World Health Organization, 2023). Pf sporozoites are transmitted to humans during the bite of infected *Anopheles* mosquitoes. After deposition in the skin, the sporozoites enter blood capillaries to access the liver and infect hepatocytes for their further differentiation and multiplication. Antibodies against the major Pf sporozoite surface antigen, circumsporozoite protein (PfCSP), can prevent infection in humans suggesting that passive immunisation strategies based on human monoclonal anti-PfCSP antibodies could be deployed to prevent malaria disease in vulnerable populations (Gaudinski et al., 2021; Kayentao et al., 2022, 2024).

Protective anti-PfCSP antibodies target the central repeat domain and the N-terminal junction (Triller et al., 2017; Oyen et al., 2017, 2018; Kisalu et al., 2018; Tan et al., 2018; Murugan et al., 2020; Wang et al., 2020; Kratochvil et al., 2021; Martin et al., 2023; Williams et al., 2024). These domains are characterised by a large number of repeating four amino acid (aa) asparagine-alanine-asparagine-proline (NANP) motifs in the central domain and NANP motifs interspersed with a few asparagine-valine-aspartate-proline (NVDP) motifs in the N-junction, which connects the repeat domain to the N-terminus and contains a single asparagine-proline-aspartate-proline (NPDP) motif. Although most anti-repeat and anti-N-junction antibodies show some degree of cross-reactivity to NANP, NVDP and NPDP motifs, due to the strong similarity in sequence and structure, highly cross-reactive human antibodies are rare (Triller et al., 2017; Oyen et al., 2017, 2018; Kisalu et al., 2018; Tan et al., 2018; Murugan et al., 2020; Wang et al., 2020; Williams et al., 2024).

We previously reported that cross-reactivity to the PfCSP central repeat domain and N-junction is linked to antibody affinity, suggesting that the immunoglobulin (Ig) gene somatic hypermutation (SHM) process, in which B cells accumulate mutations during ongoing immune responses in germinal center (GC) reactions, might contribute to the generation of high-affinity cross-reactive antibodies (Murugan et al., 2020; Thai et al., 2023). Here, we addressed this question by experiments and molecular dynamics simulation for the recombinant monoclonal antibody (mAb) 4493, a somatically mutated, highly cross-reactive and protective human anti-PfCSP mAb (Murugan et al., 2020). By generating a panel of mAb 4493 variants with different SHM profiles, we delineated the role of the SHM process in gaining cross-reactivity and affinity. Our simulations reveal how SHM alters the flexibility of the paratope in mAb 4493. The increase in flexibility of the antibody-antigen complexes reduces the entropic penalty associated with binding to NANP repeat sequences and the N-junction, thereby enhancing the antibody’s affinity for its target epitopes. Furthermore, our data show an intricate inverse relationship between antibody affinity and *in vivo* serum stability that extends to other VH3-49+Vκ3-20 anti-CSP mAbs with high similarity to mAb 4493.

## RESULTS AND DISCUSSION

### SHMs increase mAb 4493 affinity and cross-reactivity

The somatically mutated antibody, here referred to as wild-type (wt) mAb 4493 has high affinity to peptides representing the PfCSP central repeat domain and N-junction (Murugan et al., 2020). The VH3-49 and Vκ3-20 amino acid (aa) sequences of wt mAb 4493 differ from those in its clonal relative mAb 4142 and the predicted germline (GL) precursor by four and one replacement SHM, respectively (Fig. EV1A). The VH3-49 SHMs are found at positions H.Y56N and H.T59R in complementarity-determining region 2 (CDR2) and at positions H.Y62N and H.A63P in framework region 3 (FWR3), while the single Vκ3-20 SHM K.S32T is located in CDR1 (Fig. 1A). The IgH CDR3 (HCDR3) sequence in the naive precursor remains unknown. Therefore, we considered the two aa residue differences in the HCDR3 of wt mAb 4493 in comparison to mAb 4142 to be SHMs (H.A102E; H.Y108S; Fig. EV1A and Fig. 1B). In the co-crystal structure of Fab4493 with the PfCSP N-junction and central repeat peptides, only H.Y56N interacted with the peptide backbone (residue N9 of N(A/V)(N/D)PNANPN(A/V)(N/D)P), whereas the rest of the mutations did not make direct contacts (Fig. 1A; Murugan et al., 2020).

**Figure 1.**
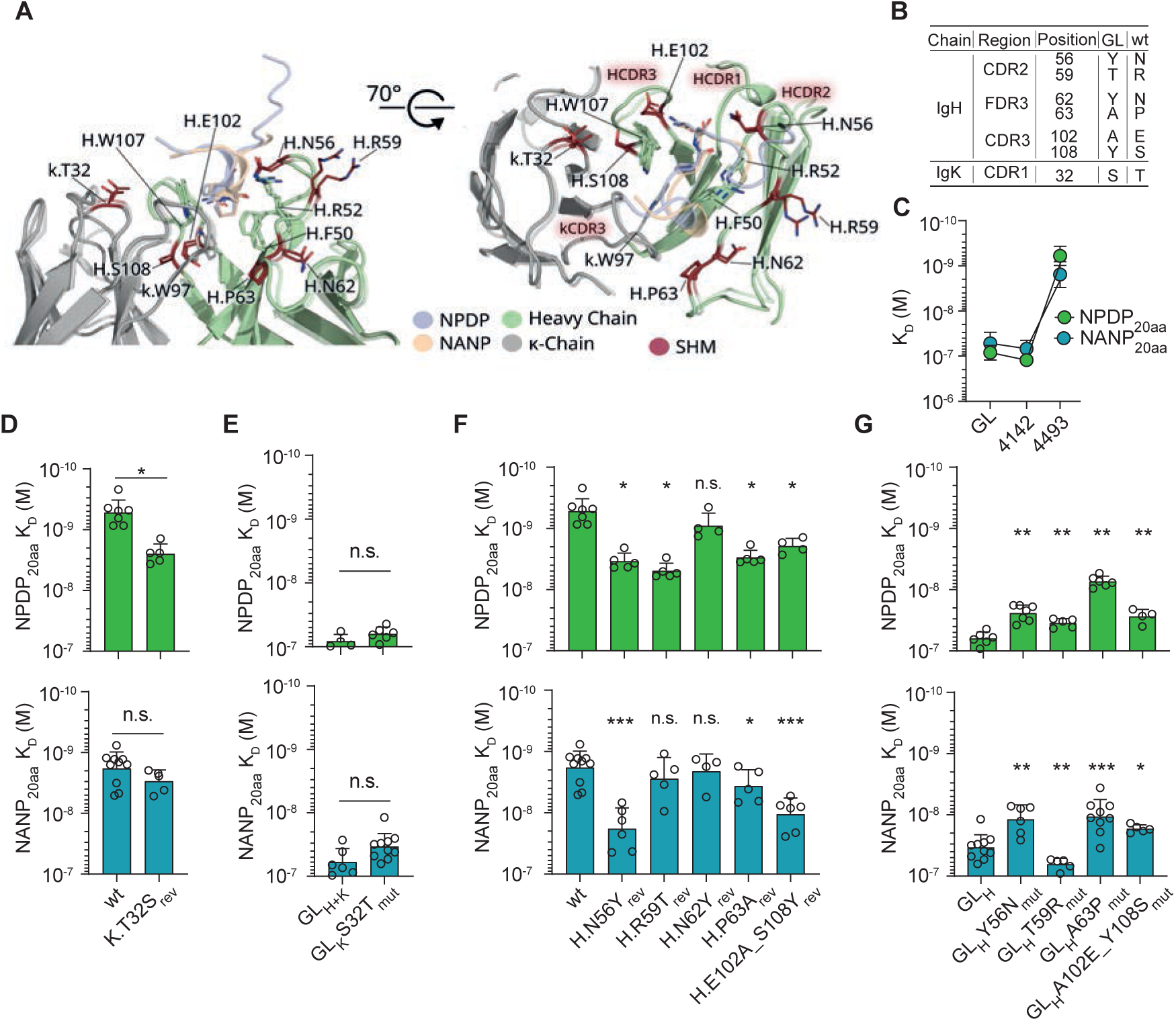
SHM-mediated affinity maturation of the PfCSP mAb 4493 to N-junction and NANP repeat epitopes. **A.** Two superimposed co-crystal structures of wt mAb 4493 bound to NPDP (blue) or NANP (orange) peptides showing the conserved positioning of the central proline-asparagine motif (N(A/V)(N/D)PNANPN(A/V)(N/D)P with P8 and N9 shown as sticks) of the peptides in the core of the paratope. The IgH and Igκ chains are shown as cartoons and the corresponding interacting residues as sticks colored i green and grey, respectively. SHMs are indicated by dark red sticks colored by atom-type (Nitrogen: Blue, Oxygen: Red). PDB IDs: 6o29, 6o2c (Murugan et al. 2020). B. Table depicting the aa replacement SHMs observed in the IgH and Igκ of wt mAb 4493. C. Affinity (equilibrium dissociation constant) of wt mAb 4493, mAb 4142, and the mAb 4493 GL version (GL_H+K_) to PfCSP junction (NPDP_20aa_: NPDPNANPNVDPNANPNVDP) and central repeat (NANP_20aa_: NANPNANPNANPNANPNANP) peptides measured by SPR. D-G. NPDP_20aa_ (top) and NANP_20aa_ (bottom) affinities of mAb 4493 (wt), 4493 GL_H+K_, and 4493 variants with GL reversions (rev; D, F) or mutations (mut; E, G) at the indicated positions in their Igκ (D, E) and IgH (F, G) chains. GL_H_ and GL_K_ indicate GL IgH and Igκ, respectively. Bars represent geometric means from at least four independent measurements (circles) and error bars are computed from geometric standard deviation factors (C-G). *P < 0.05, **P < 0.01, ***P < 0.001, n.s. = not significant, two-tailed Mann-Whitney test compared to either wt (D, F), GL_H+K_ (E) or GL_H_ (G).

To better understand the impact of the SHM process on wt mAb 4493 binding to the peptides, we generated a germline version of the antibody by reverting all SHMs in the IgH and Igκ chain (GL_H+L_) and determined the affinity to two 20 aa long peptides representing the N-junction (NPDP_20aa_: NPDPNANPNVDPNANPNVDP) and central repeat (NANP_20aa_: NANPNANPNANPNANPNANP) by surface plasmon resonance (SPR). Compared to mAbs GL_H+L_ and 4142 with similar binding strengths and reactivity profiles, the wt mAb showed 140-fold and 33-fold higher affinities to the NPDP_20aa_ and NANP_20aa_ peptides, respectively, suggesting that mAb 4493 affinity matured predominantly towards the N-junction and less to the central repeat (Fig. 1C, DATASET EV1).

To understand the roles of the individual mutations, we first reverted single SHMs in the wt antibody to the corresponding GL aa and assessed potential changes in affinity. For cases with a loss in affinity, we confirmed the role of the mutation by introducing the single SHMs in the reverted GL version (DATASET EV1). All IgH and Igκ mutations were assessed as single aa exchanges, while the HCDR3 mutations were tested in combination. Compared to wt mAb 4493, reversion variant K.T32S_rev_ lost affinity to the NPDP_20aa_ but not strongly to the NANP_20aa_ peptide, whereas introducing K.S32T in mAb GL_H+K_ (GL_H_K.S32T_mut_) did not result in any significant affinity differences (Fig. 1D and E). Thus, the K.S32T mutation increased the NPDP_20aa_ affinity only in the context of the other IgH SHMs. Similarly, all IgH SHM reversions (H.Y56N, H.A63P and H.A102E_Y108S) except H.N62Y reduced the affinity to both peptides compared to the wt antibody (Fig. 1F), whereas H.R59T reduced the affinity to NPDP_20aa_ only. Accordingly, introducing these mutations into the GL antibody, increased the affinity to the respective peptides (Fig. 1G). The overall stronger role of the SHM process in binding to the N-junction compared to the repeat were reflected by changes in the peptide association (k_on_) and/or the dissociation (k_off_) rates, with a more pronounced impact on binding to the NPDP_20aa_ (increased k_on_ in GL_H_Y56N; decreased k_off_ in GL_H_T59R, GL_H_A56P and GL_H_A102E_Y108S; Fig. EV1B) than the NANP_20aa_ peptide (decreased k_off_ in GL_H_Y56N; Fig. EV1C; DATASET EV2). Notably, the H.Y56N mutant, in which the asparagine could form hydrogen bonds with the peptide backbone, increased affinity to both peptides by distinctly modifying the on-and off-rates of the NPDP_20aa_ and NANP_20aa_ peptides, respectively. In summary, by reducing the kinetic off-rates the SHMs in wt mAb 4493 improved the affinity to NPDP_20aa_ more so than to NANP_20aa_.

### The SHMs in wt mAb 4493 impact cross-reactivity non-additively

To determine whether the effects of the SHMs on affinity were independent of each other, we focused on H.Y56N, H.T59R and H.A63P, the three VH mutations with a significant increase in affinity to at least one of the peptides compared to the GL_H_ (Fig. 1G). We generated antibody variants with these three mutations individually and in specific combinations and measured affinity to the NPDP_20aa_ and NANP_20aa_ peptides (Fig. 2). To determine the gain in affinity, we calculated free energy values (ΔG) from the equilibrium dissociation constant values (K_D_) of all antibody variants and computed the difference (ΔΔG) by subtracting the free energy of the antibody without VH SHMs (GL_H_). The three SHMs in all combinations mediated increased binding with more negative ΔΔG values, when measured for binding to NPDP_20aa_ than to NANP_20aa_. For NPDP_20aa_, the experimentally measured values (Observed) of the combined mutants were similar to the additive values of the corresponding individual mutants (Expected), suggesting that the mutations largely improved affinity independently of each other (Fig. 2A). In contrast, synergistic binding to NANP_20aa_ was observed for H.T59R in combination with H.A63P but not with H.Y56N (Fig. 2B). Moreover, the experimentally observed ΔΔG values of the H.Y56N, H.T59R and H.A63P triple mutant antibody were more negative than the expected values based on the measured affinities of the individual mutants, corresponding to a non-additive gain in affinity to NANP_20aa_.

**Figure 2.**
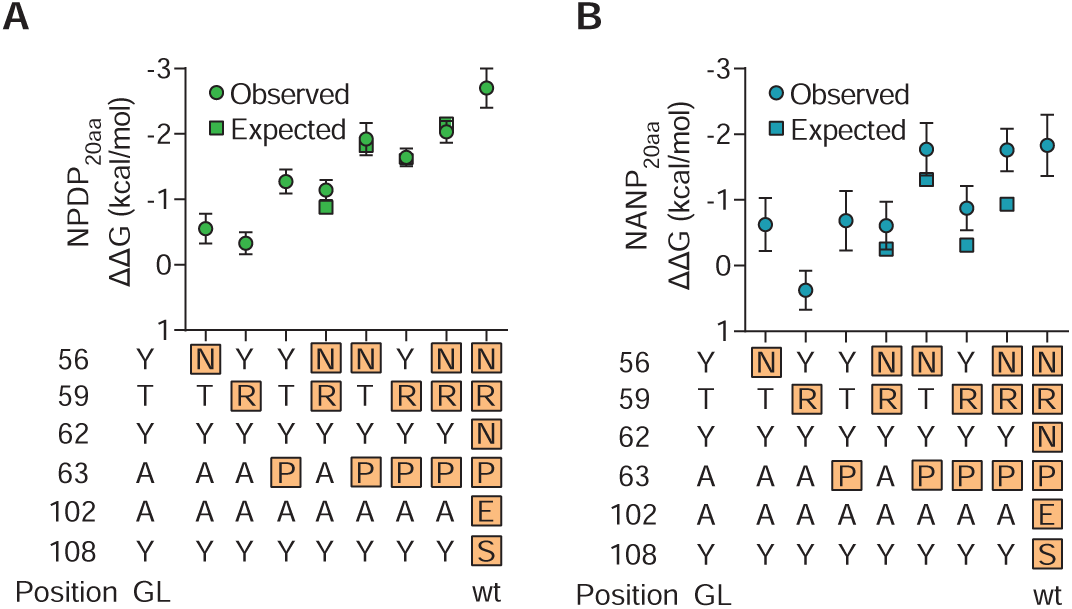
SHMs H.Y56N and H.A63P cooperatively enhance the affinity of mAb 4493 GL_H_ to the PfCSP central repeat, but not the N-junction. A, B. Binding free energy difference for NPDP_20aa_ (A) and NANP_20aa_ (B) between mAb 4493 GL_H_ and mAb 4493 GL_H_ variants with VH SHMs at the indicated positions (highlighted). Filled circles indicate mean and SEM of at least four independent experimental measurements (Observed). Binding free energy differences calculated assuming that the effects of the mutations are additive are shown as filled squares for reference (Expected).

Correspondingly, for the NPDP_20aa_ peptide, the improvement in binding kinetic rates of the individual SHMs was retained when they were combined with other SHMs (Fig. EV2A). In contrast, the kinetic binding parameters to the NANP_20aa_ peptide showed non-additive behaviour, with a decrease in off-rate in the presence of H.Y56N and H.A63P mutations (Fig. EV2B).

To better understand the generalizability of these findings, we studied two additional VH3-49+Vκ3-20 mAbs that carried some of the SHMs that we observed in wt mAb 4493 (Murugan et al., 2020; Oludada et al., 2023). mAb 3686 was isolated from the same individual as wt mAb 4493 after repeated exposure to live sporozoites under chemoprophylaxis (Mordmüller et al., 2017), whereas mAb 0786 was isolated from an individual immunized with radiation-attenuated sporozoites (Mordmüller et al., 2022; Oludada et al., 2023). Instead of Arg as in wt mAb 4493, the Thr at position H.59 was mutated to Ala and Ser in mAb 3686 and mAb 0786, respectively, whereas the H.Y56N mutation was present in both mAbs and Ala at H.63 remained unmutated (Fig. EV2C and D). Introducing Arg at H.59 in either mAb did not increase their affinity, whereas reverting H.N56 to the GL Tyr and introducing Pro at H.63 significantly decreased and increased the affinity to both peptides, respectively (Fig. EV2 E and F). Thus, the H.Y56N and H.A63P mutations strongly affected the affinity and cross-reactivity of all three VH3-49+Vκ3-20 mAbs, indicative of convergent affinity maturation trajectories.

### SHMs reduces the entropic cost of antigen binding by altering paratope flexibility

Our experimental observations show the strong impact of SHMs, such as H.A63P, in improving antibody affinity and cross-reactivity, despite the lack of evidence for direct interactions between these residues with the peptide antigen in the mAb 4493 crystal structures (Murugan et al., 2020). To investigate whether these mutations influenced the antibody’s interaction with the flexible peptides through transient contacts that were not resolved by X-ray crystallography, we carried out conventional Molecular Dynamics (MD) simulations of 0.5 μs duration of the holo (with bound peptide) and apo (without bound peptide) wt and GL_H+K_ antibodies (Fig. 3).

**Figure 3:**
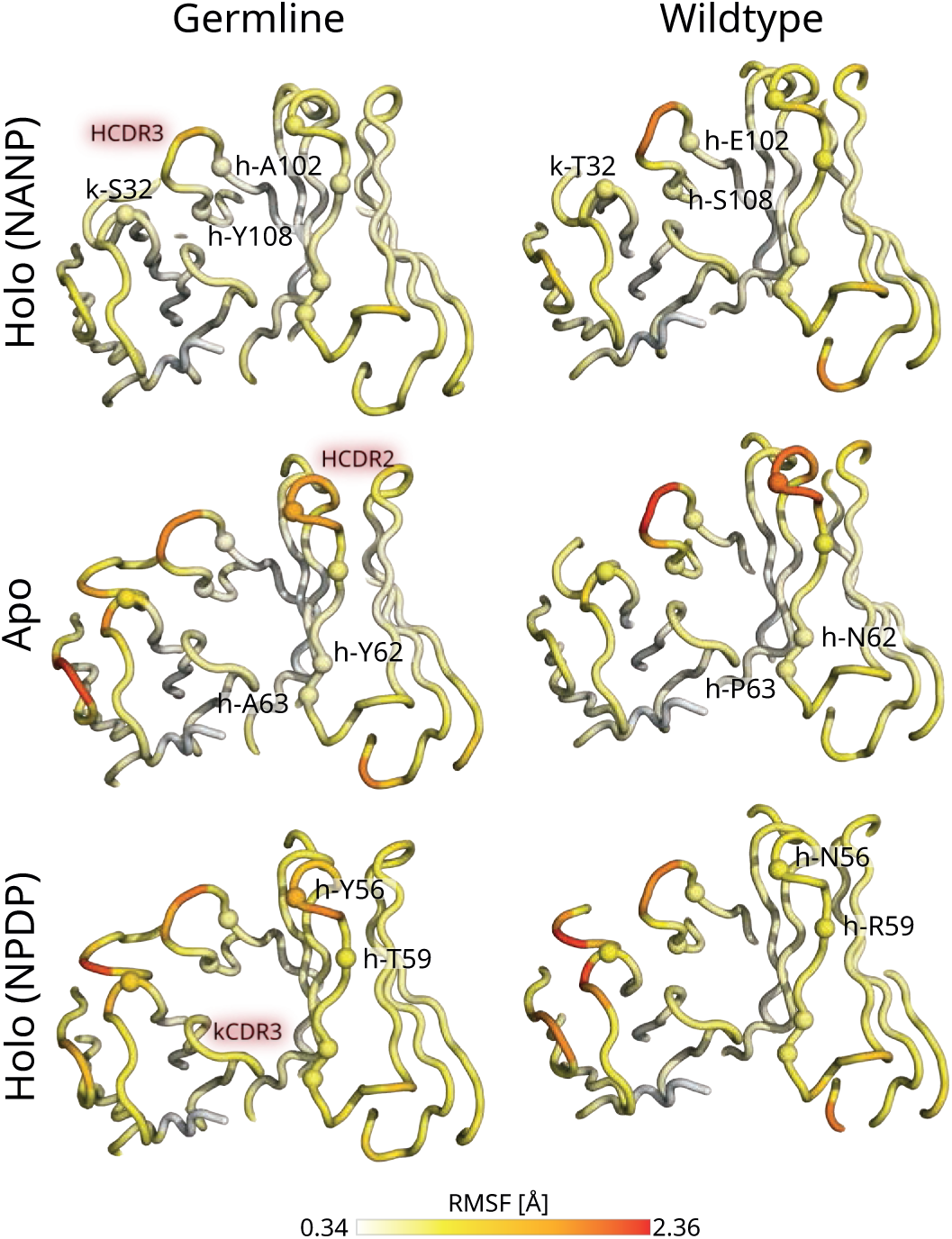
SHMs redistribute the dynamic fluctuations of the mAb4493 antibody over its structure. While the apo-mAb4493 overall becomes less flexible, the holo-mAb4493 states increase their flexibility (Table. 1). During molecular dynamics simulations, the fluctuations of the apo-mAb4493 are higher in HCDR2 and HCDR3 and lower in the Igκ chain of the wt mAb 4493 compared to the GL_H+K_ variant. In the NANP-complex, HCDR3, and in the NPDP-complex, the Igκ chain, display increased fluctuations in the wt mAb 4493 compared to the GL_H+K_ variant. Ribbon representations of the antibodies are colored by RMSF and mutations are indicated by spheres. Residues and relevant CDR loops are labelled only once for visual clarity. RMSF values were calculated for Cα atoms after alignment of the coordinates from trajectory frames to those in the first frame.

The SHMs in the IgH CDRs replace GL_H+K_ residues with smaller and less hydrophobic residues (H.Y56N, H.Y102S and H.A108E) resulting in an overall increased mobility of wt HCDR2 and HCDR3, compared to their GL_H+K_ counterparts in the apo-state. In contrast, the RMSFs of the Igκ residues are decreased in the apo-state, likely due to the K.S32T mutation improving interactions between KCDR1 and KCDR2.

Estimation of the entropies of the holo- and apo-states of wt mAb 4493 compared to the GL_H+K_ antibody using a coarse-grained quasi-harmonic model shows that the flexibility of the apo-antibody is overall lower than that of the holo-states, resulting in a lower entropic penalty for peptide binding of the wt compared to the GL_H+K_ antibody (Table 1). In contrast, binding enthalpies computed with the MMPBSA method (including a solvation free energy term) (Genheden & Ryde, 2015a; Valdés-Tresanco et al., 2021) were favourable but showed no significant difference between GL_H+K_ and wt antibody 4493.

The data indicate an overall reduction of the entropic cost of binding to NPDP and NANP peptides upon maturation from the GL_H+K_ to the wt antibody due to changes in antibody flexibility in the holo and apo states. Although the entropy estimation presented here is of a qualitative nature and does not provide complete entropic contributions, the computed changes are consistent with the experimentally determined K_D_ values (Fig. 1C, Table. 1).

**Table 1:** SHMs decrease the entropic cost without affecting the enthalpy for binding of mAb 4493 to epitope peptides*.

| Quasi Harmonic Entropy estimation | | | $\Delta S/\Delta H$ estimation | | |
| --- | --- | --- | --- | --- | --- |
| Antibody | Peptide | $-T\langle S \rangle$ [kcal/mol] | | NANP [kcal/mol] | NPDP [kcal/mol] |
| Wildtype (WT) | NPDP | $-717.91 \pm 0.22$ ● | Entropic penalty of binding<br>$-T\Delta S_{\text{Peptide}}^{\text{Antibody}} = -T \langle S_{\text{Complex}} \rangle - \langle S_{\text{Antibody}} \rangle - \langle S_{\text{Peptide}} \rangle$ | | |
| | NANP | $-703.87 \pm 0.17$ ● | | | |
| | w/o peptide | $-633.97 \pm 0.15$ ● | | | |
| Germline (GL) | NPDP | $-701.30 \pm 0.19$ ● | $-T\Delta S^{\text{GL}}$ | $90.30 \pm 0.30$ | $73.38 \pm 0.31$ ● |
| | NANP | $-686.27 \pm 0.16$ ● | $-T\Delta S^{\text{WT}}$ | $64.32 \pm 0.26$ | $48.39 \pm 0.30$ ● |
| | w/o peptide | $-642.35 \pm 0.21$ ● | $\Delta -T\Delta S^{\text{GL-WT}}$ | $25.98 \pm 0.40$ ~ | $24.99 \pm 0.43$ |
| w/o 4493 | NPDP | $-134.22 \pm 0.13$ ● | Binding enthalpy (MMPBSA) | | |
| | NANP | $-132.34 \pm 0.12$ ● | $\Delta E^{\text{GL}}$ | $-47.00 \pm 28.29$ ~ | $-50.56 \pm 20.47$ |
| | | | $\Delta E^{\text{WT}}$ | $-45.66 \pm 11.39$ ~ | $-53.20 \pm 42.26$ |
| | | | $\Delta \Delta E^{\text{GL-WT}}$ | $-1.34 \pm 30.50$ ~ | $2.64 \pm 46.96$ |

\*Entropies were calculated from the last 60 ns of the all-atom simulation replicate trajectories by coarse-grained (C-alpha atom) quasi-harmonic entropy estimation. Mean and standard deviation values were computed by bootstrapping over the analysed trajectory frames. Estimates for wt mAb 4493 and the GL<sub>H+K</sub> precursor were from independent simulations with the corresponding peptide, and values for without antibody (w/o 4493) were from independent simulations of the peptides only. Binding entropy and enthalpy of wt and GL<sub>H+K</sub> antibodies for the NANP and NPDP peptides. Enthalpies were computed as averages with the MMPBSA model using all-atom structures across the last 60 ns of antibody-peptide complex trajectories.

Interaction footprint analysis of the GL_H+K_ and wt holo-antibody simulations showed no formation of additional persistent peptide-antibody contact points compared to the crystal structures. While the H.Y56N mutation contributes to the persistent contacts between the paratope and epitope, none of the other SHMs form any persistent contacts with the epitope. Most SHMs are located in the proximity of the constant contact points (core paratope: H.F50, H.R52, H.N56, H.I101, H.W107,and K.Y33, K.W96) and form transient contacts (peripheral paratope) (Fig. 4). The H.T59R and H.A63P mutations allow for transient contacts between the NPDP peptide and the germline residue H.K67 in the wt antibody. Similarly, K.S32T improves transient contacts of the NANP peptide with the Igκ chain peripheral paratope (KCDR1).

**Figure 4:**
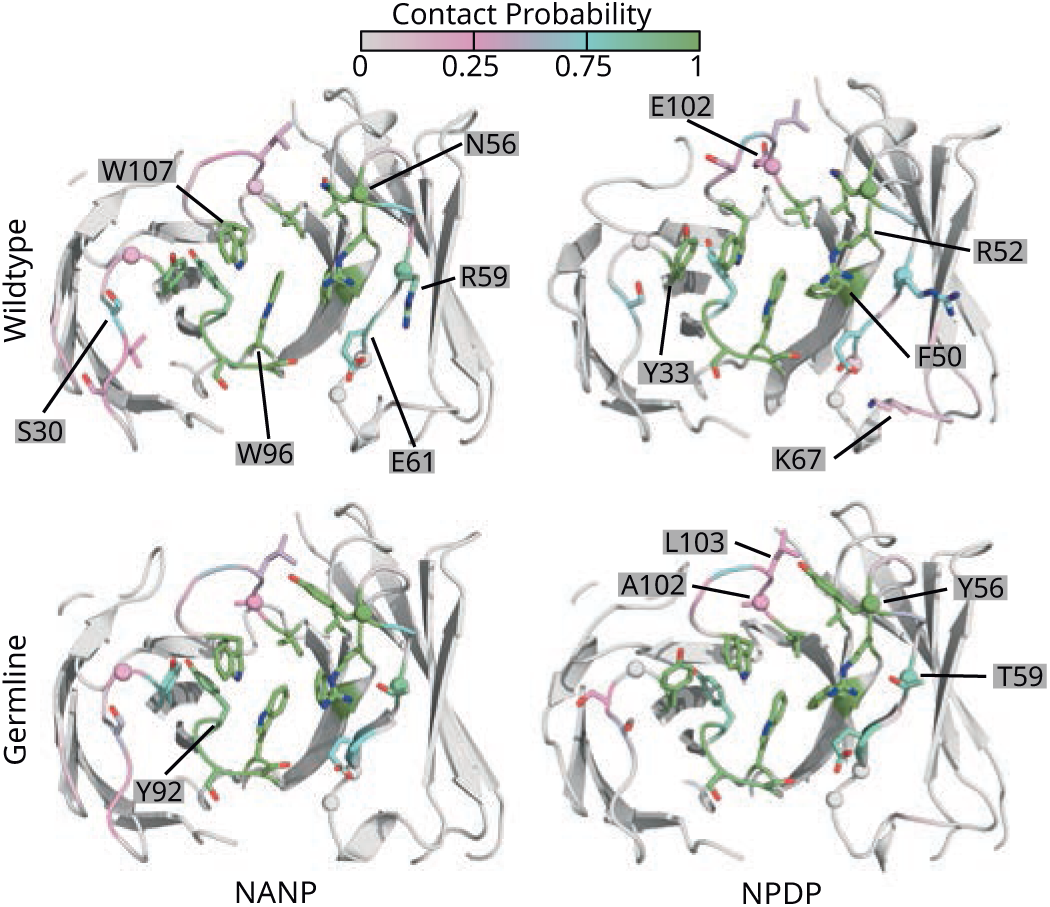
Maturation does not affect contact probabilities in the core paratope. Cartoon representations of the antibodies coloured by contact probability (#frames with contact / #number of conventional MD frames analysed). Peptides are omitted for clarity. Residues with a contact probability greater than 0.1 are shown as sticks coloured by atom type, while the positions of somatic hypermutations are indicated by spheres. For clarity, relevant contacting residues are only labelled once in the figure. The core paratope consists of residuesH.F50, H.R52, H.N56, H.I101, H.W107, and K.Y33, K.W96.

To summarize, mAb 4493 affinity maturation reduces the entropic cost of binding for both peptides through selected SHMs in the peripheral paratope without affecting the enthalpy of epitope binding and without the formation of additional contacts.

### Epitope affinity and *in vivo* serum stability of mAb 4493 are inversely related

We previously reported that the serum concentration of mAb 4493 upon passive transfer is at least 2-fold lower compared to other mAbs (Murugan et al., 2020), limiting its value for therapeutic applications. To improve its stability, we generated mAb 4493 variants by evaluating the presence of computationally derived stability violations and potential post translation modification sites using the empirical Multi-Attribute Method (MAM; (Rogers et al., 2015)).

To assess the *in vivo* stability, 100 µg of wt mAb 4493 and twelve combinatorial variants (V01-V12) were passively transferred in C57BL/6 mice and the serum mAb concentration was measured 16 h later (Fig. 5A). All but one variant (V04) showed significantly higher concentrations than the wt mAb 4493. However, the variants with increased serum stability suffered from reduced affinity to the NPDPNANPNVDPNANP and NPNANPNANPNA peptides, suggesting a link between mAb affinity and stability (Fig. 5B; DATASET EV3). To assess the protective capacity of the variants, mice passively transferred with these mAbs were challenged with *P. berghei* transgenic sporozoites expressing PfCSP and the infection burden in the liver was assessed 42h later. Antibody variants with increased serum stability compared to the wt mAb 4493 failed to enhance *in vivo* protective capacity, reflecting the cost of reduction in affinity (Fig. 5C).

**Figure 5:**
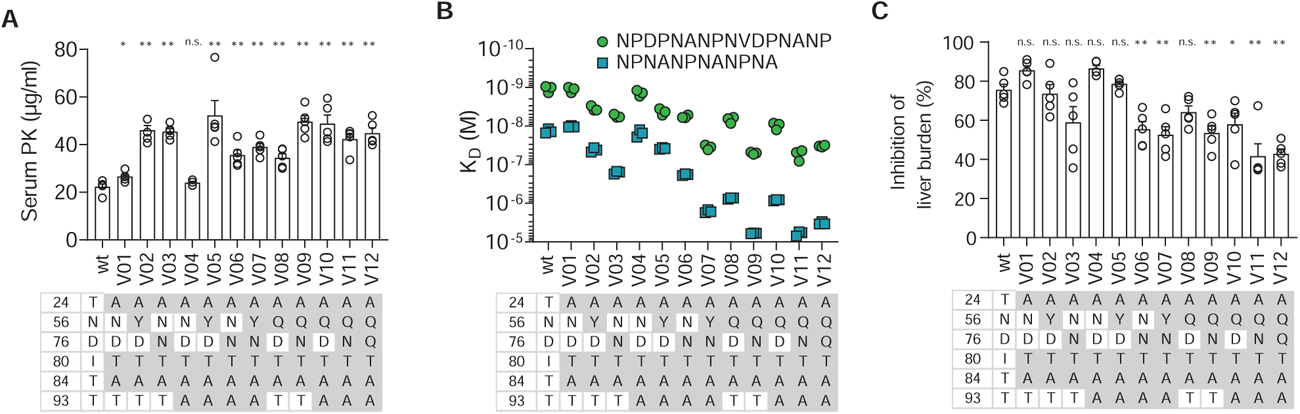
Mutations improve *in vivo* stability of mAb 4493 at the cost of affinity and protective capacity. **A-C**. Wt mAb 4493 compared to mAb variants (V01-V12) with specific mutations introduced in the IgH chain at the indicated positions (grey) for serum PK after 16 h post intravenous passive transfer of 100 µg of mAb in mice (A), affinities to NPDPNANPNVDPNANP and NPNANPNANPNA peptides measured by SPR (B), and ability to inhibit transgenic PbPfCSP sporozoites when challenged in an *in vivo* mouse model (C). Bars represent arithmetic mean and standard error of the mean (A and C). Filled circle indicates individual measurement (B). *P < 0.05, **P < 0.01, n.s. = not significant, two-tailed Mann-Whitney test compared to wt (A and C).

Similarly, the affinity-improving mutation H.A63P in mAb 0786 reduced its *in vivo* serum stability and did not enhance the mAb capacity to inhibit liver infection compared to the wt 0786 mAb (Fig. S1). The data highlight the negative association between affinity and serum stability of these VH3-49+Vκ3-20 mAbs through selective SHM.

### Maturation restricts the available conformational space of the bound peptides

To understand the impact of affinity maturation on the flexible nature of the bound epitopes, the peptide dynamics in the complexes with the wt mAb 4493 and the GL_H+K_ variant were compared in the conventional MD simulations. These simulations show that the N- and the C-terminal residues in the bound NANP and NPDP peptides remain mobile without forming persistent or transient interactions with the antibody (Fig. 4 and 6). However, binding to the antibody paratope reduces the conformational flexibility of epitope peptides (Kucharska et al., 2020) (Fig. S2, Fig. S3), which is increased upon antibody affinity maturation in the wt mAb (Fig. S2).

**Figure 6:**
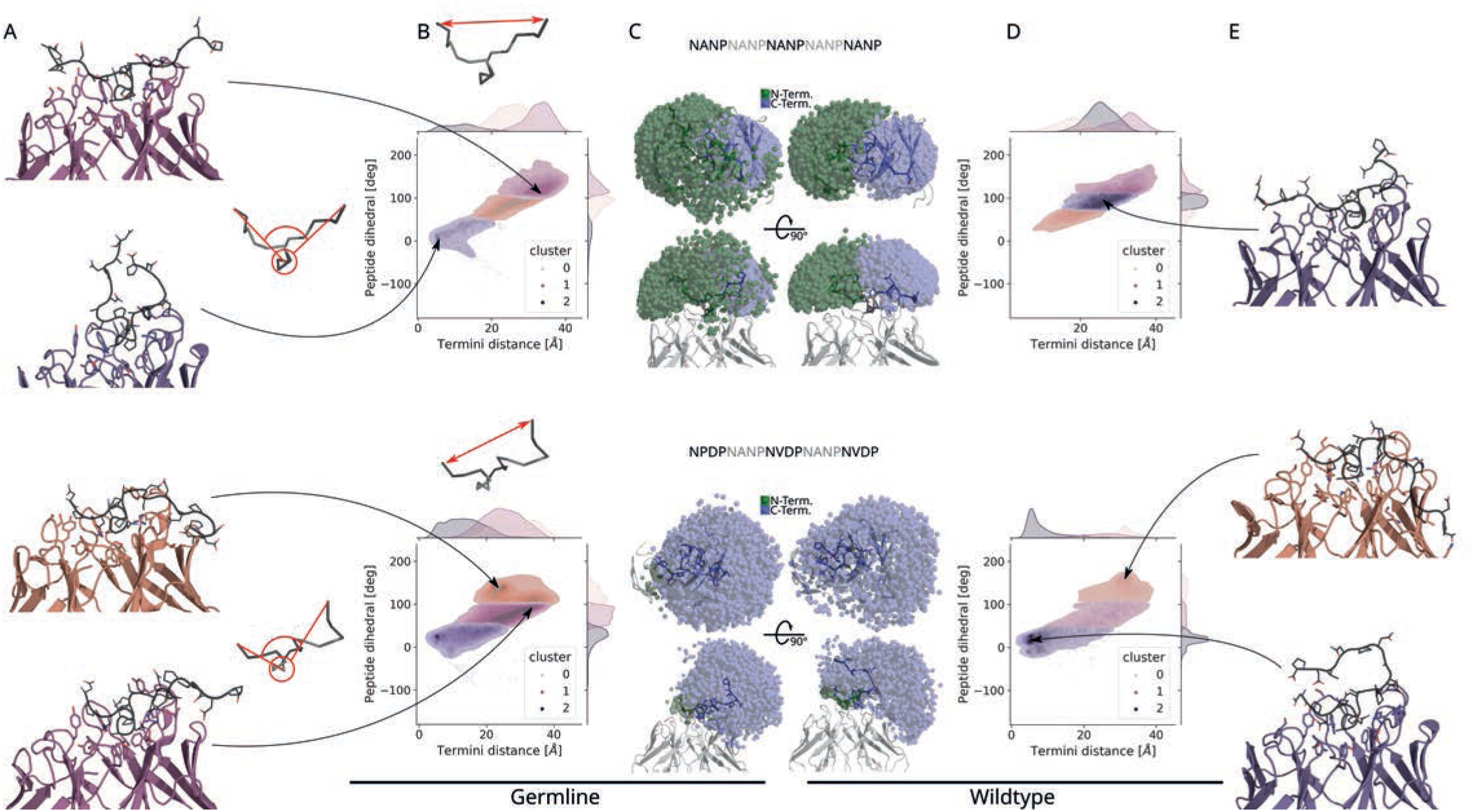
The conformational space of the NANP and NPDP peptides bound in the antibody paratope is restricted by antibody maturation. (A-E) Representative structures in the regions of highest conformational density (B, D) for the GL_H+K_ (A, B) and wt (D, E) antibodies. The antibody is shown as cartoon colored by cluster and the peptide is shown as black cartoon with side chains as sticks. Contact residues on the antibody are shown as sticks colored by atom-type (for details, refer to Fig 4). (B, D) 2D and 1D density plots show distribution of conformations sampled in conventional molecular dynamics simulations, colored by cluster number in the space described by the distance between peptide termini and the peptide dihedral angle, (indicated in red, see also Fig EV3). (C) Positions of N- and C-termini of peptides at frames extracted during the simulations are shown by green and blue spheres, respectively, from two views with GL_H+K_ shown on the left and wt shown on the right. Peptides are shown as ribbons coloured by repeat, with side chains as sticks colored by atom-type (Nitrogen: blue; Oxygen: red). Antibody depicted as gray cartoon.

Overall, the selected SHMs shifted the available conformational space to the terminal peptide repeats (Fig. 6). In detail, SHMs H.Y56N, H.T59R, H.Y62N, H.A63P, K.S32T, H.A102E, and H.Y108S restrained the NANP peptide’s N-terminus to a conformational space above the antibody’s Igκ chain, where it is prevented from making interactions with the IgH chain as in the GL_H+K_ mAb 4493. Also, maturation shifts the distribution of NANP peptide conformations from a predominantly flat, extended peptide conformation (long inter-termini distance and dihedral angle ∼ 180^◦^) in the GL_H+K_ variant to more U-shaped conformations in the wt antibody. On the other hand, antibody maturation shifts the distribution of NPDP peptide conformations towards an interaction of the peptide termini via their charges in the wt antibody. Additionally, a conformation involving interactions of the C-terminal peptide tail with FWR3 is observed in the wt antibody (Fig. 6E). It should be noted that neutralisation of the terminal charges reduces the population of conformations of the NPDP peptide with interacting peptide termini (Fig. EV4) and the positioning of the NANP peptide in the paratope can also influence the conformational distribution of the peptide (Supplementary Methods, Fig. EV4 and EV5).

In summary, we did not observe additional transient contacts between the epitope peptides and paratope in simulations compared to X-ray crystallography. However, the conformational space available to the peptides is restricted to a more U-shaped form when bound to the antibodies. This effect is increased in the wt mAb compared to the GL_H+K_ variant.

### Maturation strengthens epitope-paratope interactions up to the last point of contact during antibody-antigen dissociation

Our experimental data reveal different effects of the SHMs, especially Y56N, T59R and A63P (Fig. 1), on the binding kinetics of mAb 4493 to the NPDP and NANP peptides. To understand the influence of these mutations on the dissociation mechanism of the antibody-peptide complexes, random acceleration MD (RAMD) (Kokh et al., 2018; Lüdemann et al., 2000) simulations were carried out to simulate the dissociation of the peptides from the wt and GL_H+K_ antibodies. By applying an additional randomly oriented force on the peptide in these simulations, dissociation was observed in trajectories of up to 65 ns duration without introducing any bias on the direction of dissociation.

In RAMD simulations, unbinding of the peptide from the antibody results from loss of contact with the core paratope followed by diffusion out of the paratope via its periphery. Maturation shifts the last point of contact for both peptides towards HCDR2 and HCDR3 in the wt compared to the GL_H+K_ mAb 4493 (Fig. 7, C-F). This effect is more prominent for the NANP peptide. H.T59R is often the last point of contact for both peptides in the wt antibody, whereas H.T59 in the GL_H+K_ variant is an infrequent contact point for both peptides in the final stages of unbinding (Fig. 7, A, B, red in ΔGermline heatmaps). KCDR3 is contacted more by the NPDP peptide than the NANP peptide shortly before dissociation. In contrast, the contacts of the NANP peptide with the Igκ chain shortly before unbinding are reduced in the matured mAb (Fig. 7 A). Key residues in the pathway of dissociation from the paratope of the wt antibody are H.R52, H.N56, H.R59, H.S105, H.W107, K.Y33. Notably, residues in this group include SHMs (H.N56, H.R59) or are in their immediate neighbourhood (within 3.5 Å) (Figs. 4 and 7). Experimental data (Fig. EV1B and C) show that affinity maturation of mAb 4493 results predominantly in reduced dissociation rates: NPDP peptide dissociation is significantly slowed by the H.T59R, H.A63P and HCDR3 SHMs. NANP dissociation, in contrast, is only significantly slowed by H.Y56N. This difference is also reflected in the unbinding of NANP and NPDP in the RAMD simulations. The contacts of the NANP peptide with residues around H.N56 in HCDR2 during the dissociation are predominantly increased and those to HCDR3 are decreased in the wt antibody. For NPDP, the changes slowing dissociation that are mediated by the SHMs are predominantly within HCDR2 and FWR3; specifically, during dissociation, contact with H.R59 is increased in wt, and contact with H.N56 is reduced in wt. This indicates that the SHM process improves dissociation constants of epitope peptides from the paratope not through stabilisation of the bound state, but by improving transient interactions during dissociation.

**Figure 7:**
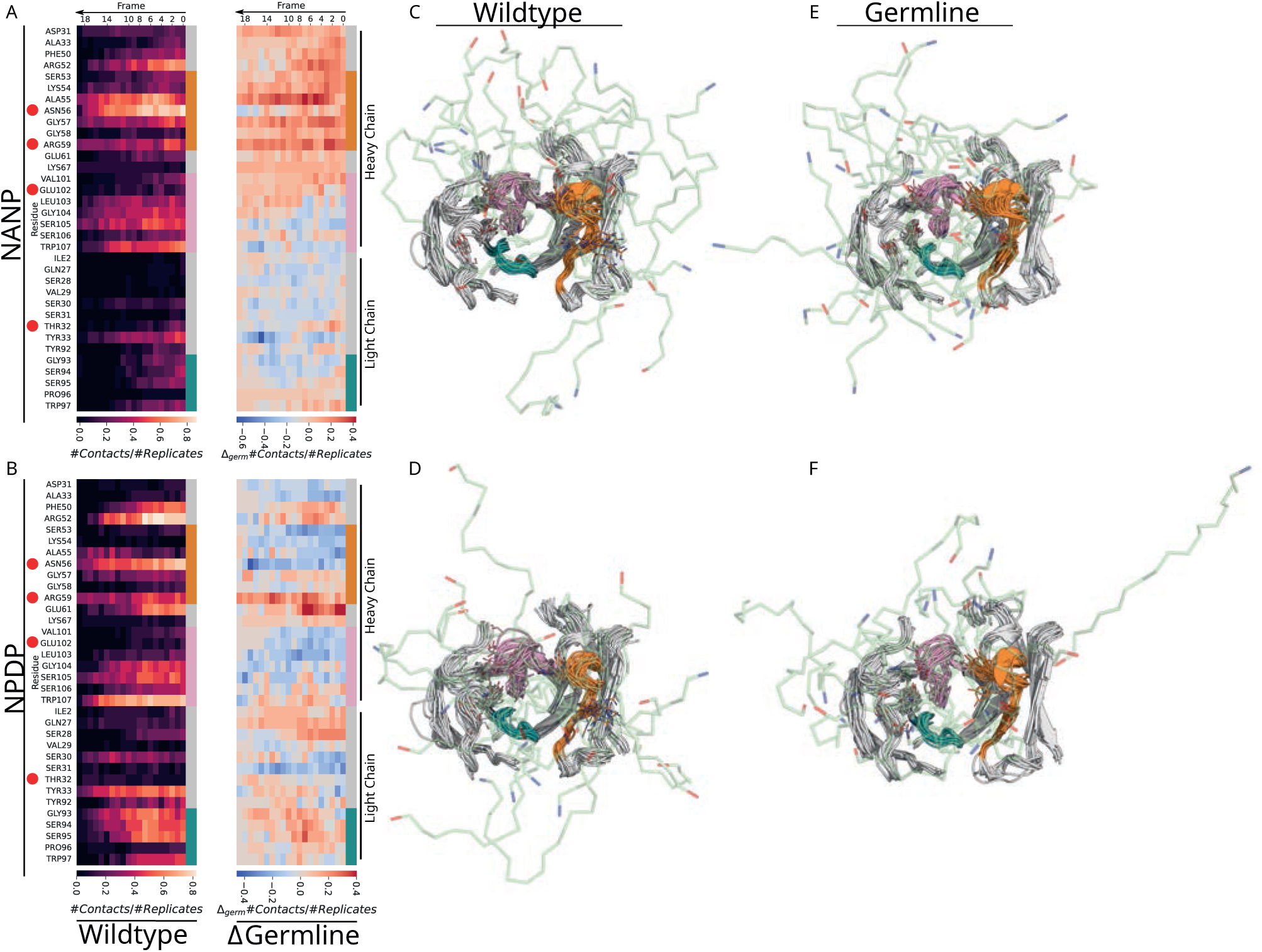
The NANP and NPDP peptide dissociation paths from the mAb 4493 paratope shift towards the periphery by antibody maturation. This effect is dominated by H.N56 and H.R59 somatic hypermutations. (A, B) Heatmaps for (A) NANP and (B) NPDP peptides depicting the contact probability in the last 200 ps of RAMD simulations before loss of contact (peptide atoms within 5Å of paratope atoms) between epitope and paratope for the wt antibody (left) and the difference to GL_H+K_ (increasing/decreasing contact probability upon SHM, blue/red) (right). The Y-axis of the heatmaps is colour-coded to indicate antibody region in accordance with the cartoon depictions. Mutations are indicated by red dots. Frames were recorded at 10 ps intervals. (C, D, E, F) Cartoon representations of the last frame before contact loss between epitope and paratope superimposed from all analysed RAMD trajectories. Antibody shown as cartoon (HCDR2 + framework - orange, HCDR3 - pink, κCDR3 - green) with last contact residues shown as sticks. The last position of the epitope peptide in each trajectory is shown as a green ribbon with blue N-terminus and red C-terminus.

## CONCLUDING DISCUSSION

Understanding of the molecular mechanisms that underlie the maturation process of high-affinity anti-PfCSP antibodies provides a basis for the design and development of potent passive or active immunization strategies against Pf. mAb 4493 and the closely related mAbs 4142, 0786, and 3686 offered a unique opportunity to determine the role of the SHM process in the development of cross-reactivity and to define the molecular features that allow the antibodies to bind to the repeat and N-junction epitopes. The fact that the SHM in this rare class of VH3-49+Vκ3-20 antibodies with a common maturation trajectory mediated a higher increase in affinity to the N-junction suggests that the underlying selection favored binding to this region. Thus, VH3-49 mAbs differ from the more common VH3-33 anti-PfCSP mAbs with dominant repeat reactivity and affinity maturation towards NANP motifs.

mAb 4142, a clonal relative of mAb 4493 with fewer SHMs, and the unmutated mAb4493 showed similar affinity to the N-junction and to the repeat, suggesting that the PfCSP cross-reactivity of these mAbs might be germline-encoded. Highly cross-reactive VH3-49 antibodies were also induced by immunization with the RTS,S/AS01, which lacks all N-junction epitopes (Williams et al., 2024). The considerable number of repeating NANP motifs in RTS,S and the lack of minor repeat and N-junction motifs suggests that these antibodies might have already shown cross-reactivity in their germline configurations. Alternatively, VH3-49 encoded antibodies induced by RTS,S might have an intrinsic propensity to gain junction cross-reactivity when undergoing affinity maturation towards the repeat motifs.

The selection process that drives affinity maturation in germinal centers leads to improved antigen binding by selecting for B cells expressing Ig mutants with improved kinetic off-rates (Roost et al., 1995). We have previously shown that NANP-reactive anti-PfCSP antibodies affinity-mature by reducing kinetic off-rates (Murugan et al., 2018, 2020). We observe a similar role for SHM in reducing the kinetic off-rates of the cross-reactive mAb 4493 when bound to NPDP_20aa_ (H.T59R, H.A63P and H.A102E_Y108S) and NANP_20aa_ (H.Y56N). However, our data also show that the affinity maturation process increased the kinetic on-rate for binding to the junction NPDP_20aa_ peptide (H.Y56N). The fact that the SHM-mediated improvements in the kinetic on- and off-rates are more pronounced for binding to NPDP_20aa_ than NANP_20aa_, likely reflects the preferred maturation of mAb 4493 to the PfCSP N-junction compared to the repeat domain.

Protein flexibility has long been known to influence antigenicity and antibody maturation (Westhof et al., 1984). Earlier studies reported that affinity maturation to conformational epitopes in structured antigens such as the HIV-1 surface envelope glycoprotein and influenza virus hemagglutinin occurs through rigidification of the antibody paratopes through SHM (Kondo et al., 2019; Schmidt et al., 2013). This overall rigidification limits paratope conformational diversity and imparts epitope specificity in addition to forming new non-covalent interactions, thereby reducing kinetic off-rates of the mature antibody to its cognate antigen (Fernández-Quintero et al., 2020; Mishra & Mariuzza, 2018). In line with these studies, our simulations of wt and GL_H+K_ mAb 4493 suggest that SHM reduces the overall flexibility of the unbound apo-antibody. In contrast, we observed increased flexibility in the bound antibody-peptide complexes of wt compared to GL_H+K_ mAb 4493, resulting in a reduction in the entropic penalty to binding. Thus, SHM induced holo-specific flexibility may be essential for high affinity cross-reactive interactions to the unstructured epitopes present in the N-junction and central repeat domains. Whether these flexible paratope-epitope interactions are unique for N-junction preferred 4493-like antibodies or a general phenomenon among anti-PfCSP antibodies is not known. However, previous reports on affinity maturation through increased paratope flexibility and overall reduced stability of affinity matured antibodies suggest alternate modes through which SHM can affect antibody paratope conformations (Maeta et al., 2023; Ovchinnikov et al., 2018; Shehata et al., 2019). The RAMD simulations of antibody-antigen dissociation highlight the key role of direct protein-peptide interactions to some of the SHMs in increasing the life-time of the mature antibody. On the other hand, the RAMD simulations also show that the GL antibody residues retaining epitope-paratope interactions until the final stages of unbinding for both peptides from the mature wt antibody are located proximal to the SHMs, which themselves do not necessarily form direct contacts with the peptides. These observations highlight the capacity of SHM to directly and indirectly influence the binding kinetics of dynamic antigen-antibody complexes.

Our data show negative effects of the affinity-improving mutations on the *in vivo* serum stability of mAbs 4493 and 0786. Introduction of stability improving mutations reduced the antibody affinity, limiting the developability of these antibodies. Both mAbs were cloned from circulating memory B cells and show signs of efficient affinity maturation. Therefore, the SHMs likely did not compromise the stability of the corresponding membrane bound Ig versions as part of the antigen receptors and had no negative effect on B cell selection during GC reactions. Reports on the negative effects of SHM on the stability of mAbs are limited (Shehata et al., 2019). Therefore, whether the negative association between SHMs and stability is restricted to specific Ig genes or VH-VL chain pairs is not known. Our data indicate that the affinity maturation process might have a negative effect on the biophysical properties of VH3-49+Vκ3-20 mAbs. Although VH3-49 anti-PfCSP antibodies induced by RTS,S vaccination also have lower serum stability than VH3-33 mAbs (Fig. S4), protein engineering efforts increased the stability of a candidate VH3-49 mAb (AB-000224; Williams et al., 2024). The ability to improve the serum stability of VH3-49 antibodies without affecting their affinity may be linked to the Ig light chain usage, as for Vλ1-40 in AB-000224, and precise affinity improving mutations.

In summary, our study highlights the role of SHMs in improving the cross-reactivity between N-junction and NANP repeat epitopes by improving binding kinetics and inducing paratope flexibility upon binding. While the findings of this study are for 20aa long epitope peptides carrying N- and C-terminal charges rather than the full-length protein antigen, our results provide valuable molecular insights into the binary binding events between the matured paratope and the corresponding epitopes. The multivalence of binding of the epitope peptides affects the experimental measurements, where the number of repeated binding motifs in the peptide determines the absolute affinity values. The effect of different positions of the two peptides on the mAbs and how they represent motifs accessibility for mAb binding on the sporozoite surface could not be fully sampled in this study, but our findings suggest that peptide positioning affects the conformational space available to NANP peptides when binding to wt Fab 4493 (Fig. EV4). The combinatorial effect of peptide positioning due to the repeating sequence on binding entropy was not considered in the presented entropy estimates. Multiple successive binding events between several 4493 mAbs and full-length PfCSP could induce a spiral tertiary structure similar to previously reported cryo-EM structures with other anti-PfCSP antibodies (Martin et al., 2023; Oyen et al., 2018). To exploit the full potential of anti-PfCSP mAbs as therapeutics, future studies should address the impact of flexible paratope-epitope interactions in the context of PfCSP on the sporozoite surface. Due to the highly unstructured nature of N-junction and NANP epitopes, studies capturing the antigen-antibody binding dynamics should be performed to gain fundamental insights into how antibody binding links to protection.

## METHODS

### Generation and expression of antibody mutants

IgH and Igκ genes were synthesized (Eurofins Genomics), cloned into IgG1 and Igκ expression vectors and recombinantly expressed in human embryonic kidney 293F (Thermo Fisher Scientific) cells (Tiller et al., 2008).

### Surface plasmon resonance

Affinity measurements using SPR were performed as described previously (Murugan et al., 2018). Briefly, the Biacore T200 (Cytiva) instrument was docked with a series S sensor chip CM5 (Cytiva). Immobilization of anti-human Fab antibodies by amine coupling was performed using a Human Fab capture kit (Cytiva), following the manufacturer’s instructions. Hepes (10 mM) with 150 mM NaCl was used as a running buffer at pH 7.4. Sample and reference flow cells were used to capture equal concentrations of sample and control antibody (mGO53, (Wardemann et al., 2003)). Upon stabilizing the flow cells with a running buffer for 20 min at 10 μl/min, NPDP_20aa_ and NANP_20aa_ peptides dissolved in the running buffer were injected at the following concentrations: 0, 0.015, 0.09, 0.55, 3.3, and 20 μM. Association and dissociation of the injected peptides were performed at 25°C for 60 and 180 s, respectively. Upon completion, the flow cells were regenerated with 10 mM glycine-HCl pH 2.1. The obtained data were fitted using a 1:1 binding model with the Biacore T200 software version 2.0. Geometric means and corresponding geometric standard deviation factors were computed using GraphPad Prism (version 9.3.1).

Affinity measurement of stability improved mAb 4493 variants carrying LS mutations with peptides were measured using the Carterra LSA SPR platform and CMD200M sensor chips (Carterra) at 25°C as described before (Thai et al., 2023). In brief, the chip was directly immobilized with 10 μg/mL of variant mAbs through amine-coupling in 10 mM Sodium Acetate at pH 4.5 and unreactive esters were quenched using 1 M ethanolamine-HCl at pH 8.5. The running buffer for immobilization and quenching was 10 mM MES at pH 5.5 with 0.01% Tween-20. Upon washing non-specifically bound IgG, 2-fold serially diluted peptide series with the starting concentration 25.6 μM and 9.41 μM for NPNANPNANPNA and NPDPNANPNVDPNANP peptides, respectively, were injected. The running buffer and sample diluent for antigen injections was 1X HBSTE buffer (10 mM HEPES pH 7.4, 150 mM NaCl, 3 mM EDTA and 0.01% Tween-20). Data were collected with 120 s of baseline step, 300 s of association step and 900 s of dissociation step and pre-processed using Kinetics (Carterra) software. TitrationAnalysis tool was used to perform fitting with 1:1 Langmuir model and derive kinetic rates and affinity values as previously reported (Li et al., 2024).

### Parasite liver burden assay

The liver burden assays were performed as previously described (Flores-Garcia et al., 2019). Briefly, *Anopheles stephensi* mosquitoes infected with transgenic *P. berghei* sporozoites expressing the *P. falciparum* CSP and luciferase were kept in an incubator at 19 °C. Sporozoites from mosquitoes were harvested at days 20-23 post-infection in HBSS-2% FBS. Mice were passively immunized in the tail vein with 100 µg of antibody per mouse and challenged 16 h later with 2x10^3^ sporozoites injected intravenously. Control mice received irrelevant or no antibodies. Forty-two hours after challenge, mice were injected intraperitoneally with 100 µl of d-luciferin (30 mg/mL), anesthetized with isoflurane and the bioluminescence was measured using an IVIS Spectrum Imager, Perkin Elmer. All procedures were performed in accordance with the recommendations in the Guide for the Care and Use of Laboratory Animals of the National Institutes of Health, under protocol number MO18H419, approved by the Animal Care and Use Committee of Johns Hopkins University.

### Mosquito bite-based *Plasmodium berghei* PbPfCSP(mCherry) mouse challenge model

*Anopheles gambiae 7b* mosquitoes, an immunocompromised transgenic mosquito line derived from the G3 laboratory strain, were used for infections with *P. berghei* sporozoites that express PfCSP and the fluorescence reporter mCherry (PbPfCSP(mCherry), (Ludwig et al., 2023). Female CD1 mice (7-12-week-old) and female C57BL/6J mice (8-week-old) were bred and housed in a pathogen-free animal facility (Experimental Animal Facility of the Max Planck Institute for Infection Biology) in accordance with the German Animal Protection Law (§8 Tierschutzgesetz) and approved by the Landesamt für Gesundheit und Soziales (LAGeSo), Berlin, Germany (project numbers 368/12 and H0335/17). Mosquitoes were fed on PbPfCSP(mCherry)-infected female CD1 mice 3 days post passage (0.1-0.8% gametocytemia) for 30-45 min and kept at 20°C 80% humidity 12/12 h day/night cycle. Infected mosquitoes were offered an additional uninfected blood meal to boost sporozoite formation at 7 days post infection (dpi), and at eleven dpi were used for C57BL/6J mice challenge experiments and live sporozoite FACS staining as described below.

Female C57BL/6J mice were passively immunized by intraperitoneal injection of 100 μg of monoclonal antibodies in 200 μl PBS. Mice were randomly assigned to separate groups. After 20 h, mice were exposed to the bites of three PbPfCSP(mCherry) salivary gland-infected mosquitoes as described elsewhere (Ludwig et al., 2023). Peripheral whole blood was collected from the submandibular vein 2-3 h after challenge to quantify serum mAbs titers by ELISA. Mice with undetectable levels of mAbs were excluded from the analysis. From days 3 to 7 and on day 10 post mosquito bite, parasitemia (mCherry-positive red blood cells/total red blood cells) was measured by flow cytometry (LSRFortessa, BD Biosciences), and confirmed by Giemsa-stained thin blood smears. All infected mice were euthanized on day 7 post mosquito bite, before the occurrence of malaria symptoms. FACS data were analyzed by FlowJo V.10.8.2 and the pre-patency period was declared on the first day when parasitemia values were above the background signal of negative mice.

### Molecular model building

The peptides used in the binding experiments are longer than those in available crystal structures. Therefore, missing peptide residues were added and somatic hypermutations (GL and mutant antibodies) removed or introduced with MODELLER (Eswar et al., 2008; Webb & Sali, 2016).

The models were based on crystal structures of antibody-peptide complexes, PDB 6o2c (NANP peptide) and PDB 6o29 (NPDP peptide) (Murugan et al., 2020), truncated with PyMoL to contain only the variable region (Ovchinnikov et al., 2018) of the antibody with N-methylamine (NME) capped C-termini on the IgH chain Missing 4-aa motif (either NANP or NPDP) were added equally on both sides to the structure from PDB 6o2c and on the C-terminal side to that from PDB 6o29 (see Fig EV5).

For each system (each antibody mutant and peptide combination), 25 models of the antibody and peptide were generated with MODELLER using 3000 iterations, 3 optimization repeats, slow MD refinement and only selecting mutated or missing residue positions of the system. The models were ranked by the sum of DOPE and GA341 scores, and the quality was assessed with ProCheck (Laskowski et al., 1993) and FoldX (Laskowski et al., 1993; Schymkowitz et al., 2005).

### Conventional MD simulations

Five of the ten best models were visually selected for their peptide conformational diversity and all-atom conventional MD simulations were run for each of them with a production time of 100 ns. Simulations were carried out using the Gromacs 2021.5 (Abraham et al., 2015) software. The CHARMM36m force field (Huang et al., 2016) was used for the protein, and the charmm36m-adjusted TIP3P model (Huang et al., 2016) was used for water molecules. Water molecules within 7 Å of the peptide in the crystal structures were included in the simulation systems, which were immersed in a periodic box of solvent. The models were protonated at pH 7.5 using ProPKA (Dolinsky et al., 2007; Olsson et al., 2011), 200mM NaCl was added, and the periodic simulation boxes were neutralized by adding five (NANP, wt and GL_H+K_) or two (NPDP, wt and GL_H+K_) chloride anions.

The solvated systems were energy minimized by steepest descent (with strong restraints on the antibody and weak restraints on the peptide) and conjugate-gradients (weak restraints on all antibody and peptide atoms). They were then equilibrated using a 1 fs time-step with gradual release of the side-chain and backbone restraints (Appendix Table S1).

In production simulations (Appendix Table S2), the temperature and pressure were held constant by Nose-Hoover temperature coupling (tau = 1 ps^-1^, 298.15 K) and isotropic Parrinello-Rahman pressure coupling (tau = 5 ps^-1^, 1 atm), respectively. During production simulations, a time-step of 2 fs was used.

Bonds to hydrogen atoms were restrained with LINCS (Hess, 2008), long-range electrostatics calculated with Particle Mesh Ewald (cut-off distance of 1.2 nm), and the Verlet cutoff scheme used for short range non-bonded interactions (cut-off distance of 1.2 nm).

Trajectories were aligned to a reference structure before calculation of root mean squared deviation (RMSD) and root mean squared fluctuation (RMSF) (for C-alpha) values with MDAnalysis (Michaud-Agrawal et al., 2011).

### Entropy and Enthalpy estimation

Entropies were estimated for complexes, apo-antibodies, and peptides with the quasi-harmonic approximation (Chang et al., 2005; Karplus & Kushick, 1981) following the protocol described by Ovchinnikov *et al*. (Ovchinnikov et al., 2018) to gain qualitative insights into the entropic costs of differences in flexibility between systems. For entropy estimation, 3000 concatenated coarse-grained frames (10 frames/ns, last 60 ns of each of the 5 conventional MD replicates) of each component aligned to their mass weighted average structure were taken from independent simulations. Coarse graining was done by reducing each residue to a single bead centred at its centre of mass.

Mass weighted covariance matrices, eigenvalues and entropies were calculated with cpptraj on these trajectories and their mass weighted average structures (Roe & Cheatham, 2013). The standard deviations of the entropy estimates were calculated by bootstrapping the trajectories 60 times, using 80% of the frames for each calculation.

Binding enthalpies were estimated for GL_H+K_ and wt mAb 4493 by MM/PBSA calculations (Genheden & Ryde, 2015b; Valdés-Tresanco et al., 2021) with ‘gmx_MMPBSA’ (Valdés-Tresanco et al., 2021) using default parameters unless specified (input files provided in supplementary material). Energies were computed for atomic-detail coordinates extracted at time intervals of 0.5 ns from the last 60 ns (600 frames) of conventional MD simulations of the antibody-peptide complexes using the single trajectory protocol. Energies were estimated at 298.15K and at an ion concentration of 200mM. Electrostatic energy terms were calculated by solving the linearized Poison-Boltzman equation with an internal dielectric constant of 1 and a solvent dielectric constant of 80 and charmm36m parameters. Solvation terms were calculated based on the solvent accessible surface area with a probe radius of 1.2 Å. Entropic terms were not considered.

### RAMD simulations

Random acceleration molecular dynamics (RAMD) simulations sample possible unbinding paths of ligand molecules from receptors on the nanosecond timescale by adding an additional randomly oriented force to the ligand (Lüdemann et al., 2000). An adjusted RAMD protocol based on the RAMD 2020-v2 gromacs implementation^1^ (Kokh et al., 2018, 2020) was used to sample the unbinding of the peptides from the receptors. From the last 60 ns of each of the five conventional MD trajectories, four antibody/peptide structures were extracted at equal time intervals (yielding 20 structures in total). The extracted structures were resolvated in 3 nm padded octahedral water boxes and charge-neutralized at 200 mM NaCl by adding ions. The solvent was re-equilibrated in short NVT and NPT simulations with high restraints on the antibody (see Tab. EV1). RAMD trajectories were then generated using the same parameters as conventional MD with a maximum simulation time of 60 ns and with velocity reassignment at the beginning of each simulation. Each epitope peptide was represented as five RAMD ligand groups, each containing four residues of the peptide and experiencing a randomly oriented force of magnitude 2.625 kcal/mol/Å. RAMD trajectories were terminated when all ligand groups were beyond 4 nm from the antibody’s center of mass. The RAMD parameters were evaluated every 0.1 ps and a new random force direction assigned if a ligand group’s center of mass had moved less than 0.0025 nm since the last RAMD evaluation. Simulation frames were written to an output trajectory every 10 ps.

### Interaction fingerprint analysis of the trajectories

Interaction fingerprint analysis of the trajectories was implemented with customized MDAnalysis (Michaud-Agrawal et al., 2011) python classes.

For conventional MD simulations, the interaction fingerprints of the peptides on the antibody were calculated for the last 60 ns of all the replicates by recording contacting antibody residues, where a contact was defined as a distance below 3.5 Å between any atom of an antibody residue and any peptide atom. Contact probabilities per residue were calculated as the percentage of all frames (3000) with a contact to the residue in question.

For RAMD simulations, interaction fingerprints were calculated up until unbinding occurred. Peptides were considered unbound when no contact within 5.5 Å between peptide and antibody was present. Contact probabilities over the last 200 ps before unbinding were calculated as the percentage of replicates that had a contact for the given residue at a given frame before unbinding.

### mAb developability analysis

The 4493 Fv sequence was analyzed in conjunction with the 4493:CSP structure 6O28 (Murugan et al., 2020) using the Just–Evotec Biologics’ Abacus™ platform. Potential developability issues such as stability violations, missing residues, free cysteines, Fv N-linked glycosylation sites, deamidation sites, isomerization sites, methionine and tryptophan oxidation sites, CDR length outliers, immunogenicity predictions, known salt-bridge stabilization positions, isoelectric point, and surface patch properties were evaluated in Abacus™ and MOE (Molecular Operating Environment (MOE), 2022.02 Chemical Computing Group ULC, 910-1010 Sherbrooke St. W., Montreal, QC H3A 2R7, 2024). Based on the information obtained, residue replacements were identified and used for the generation of combinatorial variants produced in stable pools for functional analyses. The residue positions selected for modification were evaluated structurally for PfCSP binding impact using the co-crystal structure (Murugan et al., 2020), and the combinatorial variants built by including the parental residue if its position was near the paratope.

## Supporting information

Extended View Tables 1-3

## ACKNOWLEDGEMENTS

We thank Dorien Foster, Julia Gärtner, Claudia Winter (German Cancer Research, Heidelberg) and Cornelia Kreschel, Hadenal Gordon, Manuela Andres, Daniel Eyermann and Liane Spohr (Max Planck Institute for Infection Biology, Berlin) for their assistance with experiments and the DKFZ/ EMBL/Heidelberg University Chemical Biology Core Facility, especially P. Sehr, for technical assistance. We acknowledge Dr. Sarah Mudrak, Christine Siska, Pauline Smidt, Jennifer Smith-Yuen, Deborah Hopkins, and Caren Tidwell for program management and aiding this work. We thank Dr. Jean-Philippe Julien for discussion regarding the manuscript contents. We acknowledge the support of the work by the Bill and Melinda Gates Foundation (OPP1193104 to H.W., INV008062 to R.R.K., and INV-008612 and INV-043419 to G.D.T.). This research was supported by of the Spotlight Project “Synthetic Immunology” of the Engineering Molecular Systems Flagship Initiative at Heidelberg University (subproject 5 (Engineering Antibody Evolution)). A.H. and R.C.W. would like to acknowledge the support of the Klaus-Tschira foundation and the provision of computing resources by HITS and by the state of Baden-Württemberg through bwHPC and the German Research Foundation (DFG) through grants INST 35/1134-1 FUGG and INST 35/1597-1 FUGG.

## DATA AVAILABILITY

Simulation data is available under: https://doi.org/10.5281/zenodo.11470585 and extended datasets can be made available upon request.

## AUTHOR CONTRIBUTIONS

R.M., G.C., Y.FG., designed, conducted the experiments and interpreted experimental results. G.Q.H, O.E.O., G.M., conducted experiments. A.H., designed, performed simulations, interpreted results and wrote the manuscript. K.L, R.H.C.H, S.M.D, G.D.T., F.Z., E.A.L., R.R.K interpreted the results. R.M., R.C.W., and H.W. conceived the study, interpreted the results and wrote the manuscript with input from other coauthors.

## EXPANDED VIEW FIGURES

**Figure EV1:**
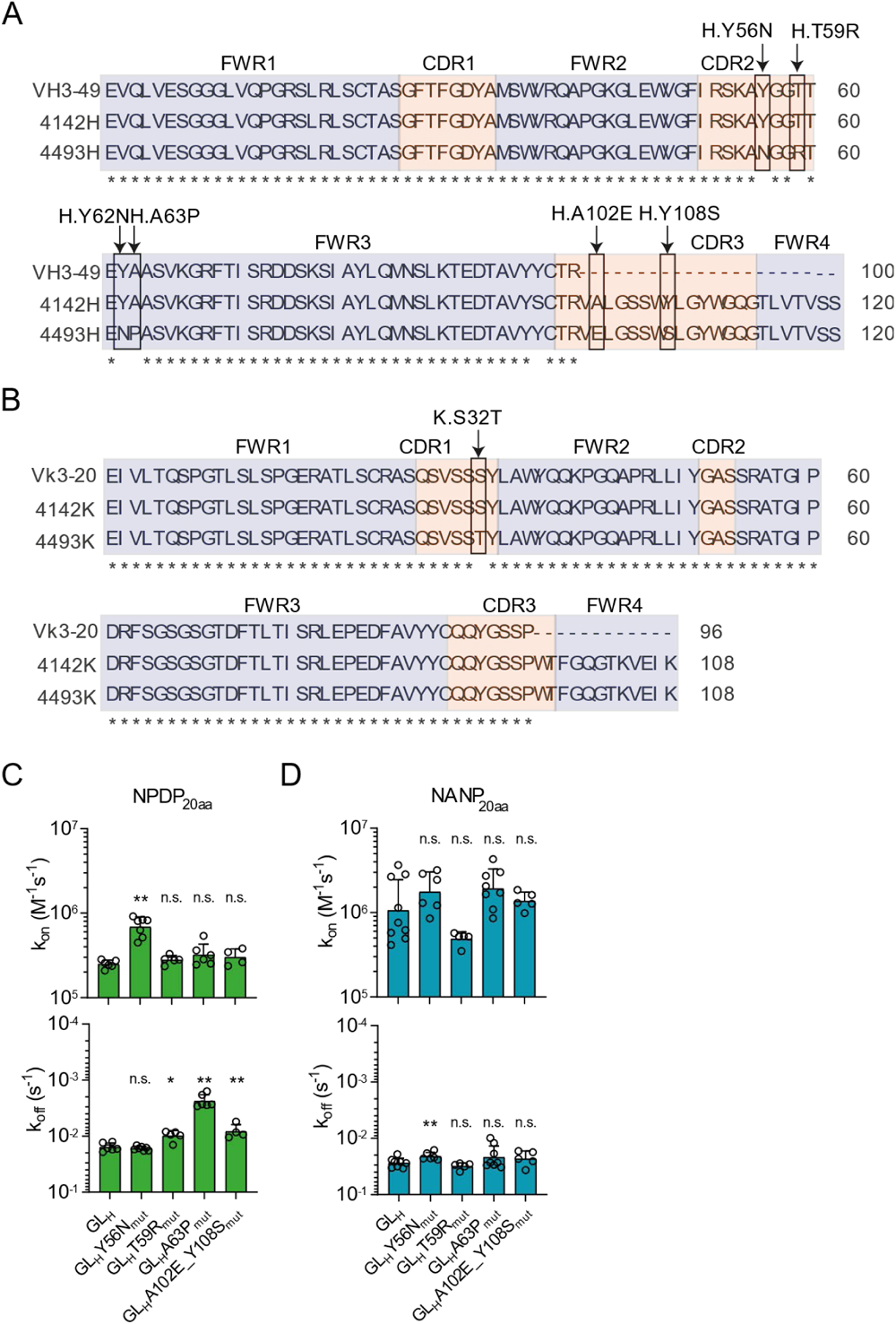
VH gene SHM increases mAb 4493 affinity to the PfCSP repeat and N-junction. **A**. Amino acid sequence alignment of the variable regions of the clonally related mAbs 4142 and 4493 to the GL VH3-49 and Vκ3-20. Framework regions (FWR) and complementary determining regions (CDR) are coloured in blue and red, respectively. Arrows indicate SHMs. **B, C**. NPDP_20aa_ (B) and NANP_20aa_ (C) kinetic on (top) and off (bottom) rates of mAb 4493 GLH and mAb 4493 GLH variants with mutations at the indicated positions in their IgH chains. Geometric mean is plotted with error bar from the geometric standard deviation factor. *P < 0.05, **P < 0.01, n.s. not significant, two-tailed Mann-Whitney test.

**Figure EV2.**
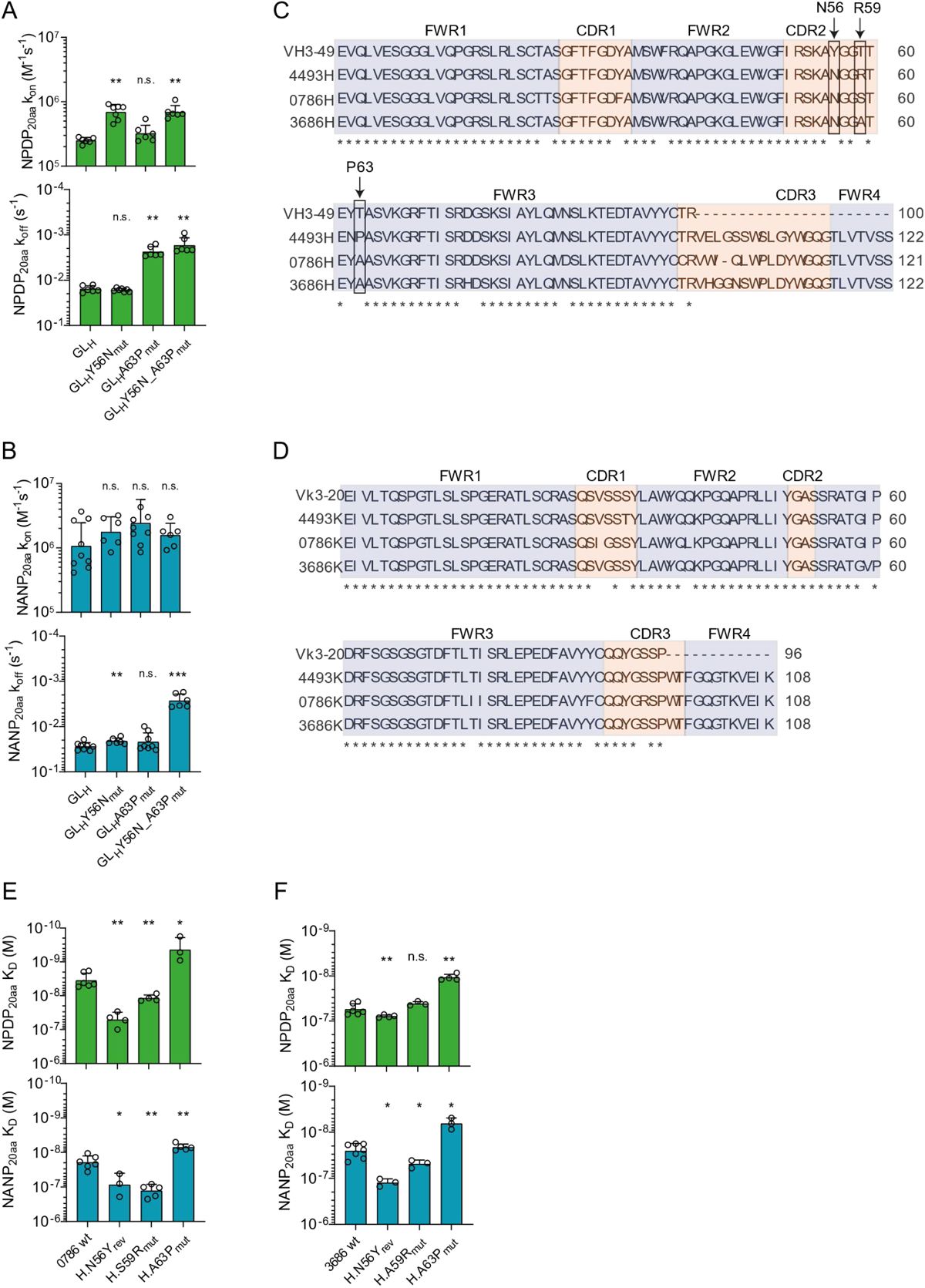
Changes in affinity and kinetic rates for SHMs in mAbs 4493, 0786 and 3686. **A, B**. NPDP_20aa_ (A) and NANP_20aa_ (B) kinetic binding on (top) and off (bottom) rates of mAb 4493 GLH and mAb 4493 GLH variants with mutations at the indicated positions in their IgH chains. **C, D**. Amino acid sequence alignment of the variable regions of the mAbs 4493, 0786 and 3686 to the GL VH3-49 (C) and Vκ3-20 (D). Framework regions (FWR) and complementary determining regions (CDR) are coloured in blue and red, respectively. Arrows indicate SHMs of interest. **E, F**. NPDP_20aa_ (top) and NANP_20aa_ (bottom) affinities of mAb 0786 wt (E) and 3686 wt (F) with either reversions (rev) or mutations (mut) at the indicated positions in their IgH chains. A, B, E and F. Geometric mean is plotted with error bar from the geometric standard deviation factor. *P < 0.05, **P < 0.01, n.s. not significant, two-tailed Mann-Whitney test.

**Figure EV3:**
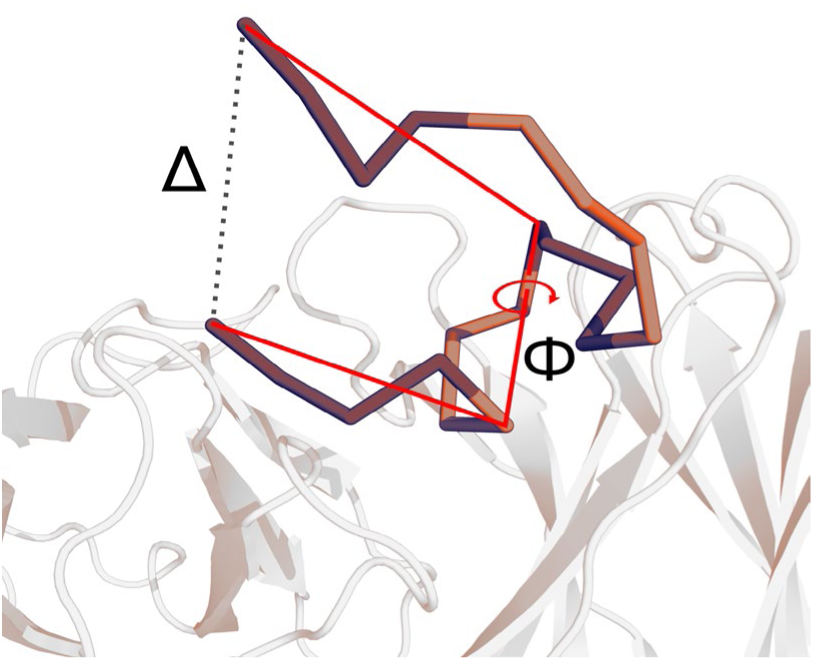
Definition of conformational descriptors of peptides monitored in conventional MD simulations and shown in Fig. 6. The antibody is shown in white cartoon, and the epitope is shown in a dark/light orange ribbon (shade changing between each consecutive peptide repeat). The dashed line indicates the distance between the epitope termini, and the red lines indicate the peptide dihedral angle (Φ) monitored.

**Figure EV4:**
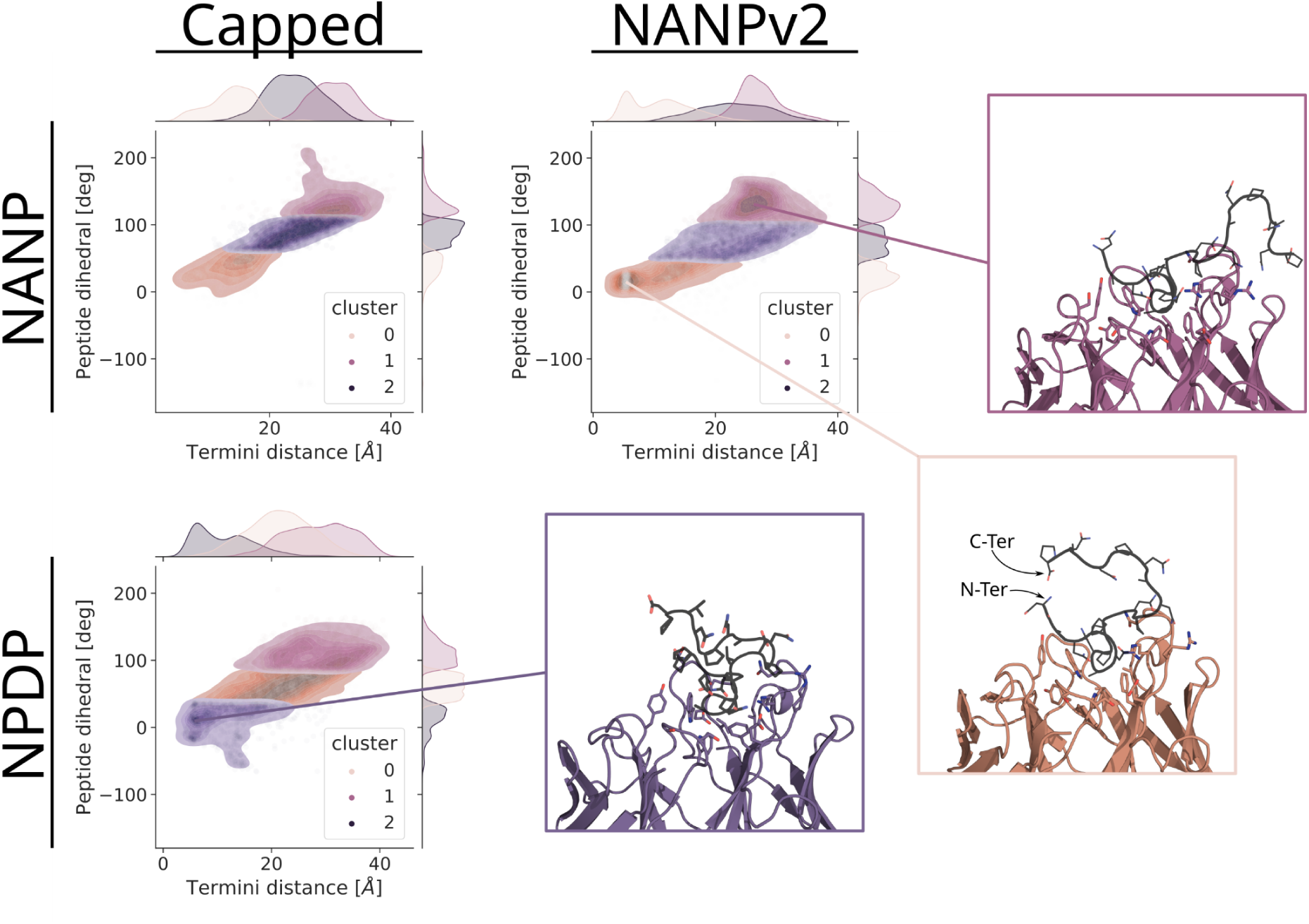
Distribution of peptide conformations in conventional MD simulations of wt antibody and capped peptides with neutral termini and of complexes with shifted NANP positioning (NANPv2, with charged termini). Scatter plots show the 2D and 1D distributions of conformations in the space of the peptide dihedral and termini distances (defined in Fig. EV3). 3D structures show representatives of high-density regions in the conformational distributions with the antibody in cartoon representation coloured by cluster. Peptides are coloured in black, with side chains shown as sticks coloured by atom type. Based on the set-up of *in vitro* experiments, the peptide termini were generated with zwitterionic charges, and this assignment was adopted for all conventional and enhanced sampling simulations. However, these charges would not be present in the full length PfCSP as the epitope peptides are located within the unordered repeat region of the protein. This could introduce differences in the binding between the biological and experimental systems. To gauge the impact of the charged termini on the interaction between antibody and epitope peptides, a set of wt mAb 4493 simulations with uncharged capped termini were performed (see Supplementary Methods). This set of simulations shows that removal of the terminal charges influences the distribution of sampled conformations in the wt antibody mainly for the NPDP peptide. Capping of the termini with neutral groups did not affect the distribution of conformations for the NANP peptide in the wt antibody compared to uncapped simulations. Conversely, removing the terminal charges from the NPDP peptide resulted in a reduced frequency of interactions between the termini. Instead, it resulted in a distribution similar to the (U-shaped) NANP peptide conformations seen in both capped and uncapped NANP wt simulations (compare Fig. 6D; and Fig. EV4, capped). Conformations with low inter-termini distances in the simulations of capped NPDP peptide in wt show no electrostatic interactions via the backbone terminal residues (Capped, NPDP). The positioning of the NANP peptide is ambiguous from the crystal structure and sequence (see Supplementary Methods). Effects of positioning on the conformational distribution were gauged by a set of simulations of the wt antibody with the NANP peptide positioned like the NPDP peptide. The NANP peptide positioning similar to the NPDP peptide resulted in a similar conformational distribution to the NPDP peptide (compare NANPv2 and Fig. 6E, NPDP wt). Conformations with interactions between the NANP peptide’s C-terminus and HCDR2 were the most frequent, and conformations with interactions between the peptide termini via their charges were the second most frequent.

**Figure EV5:**
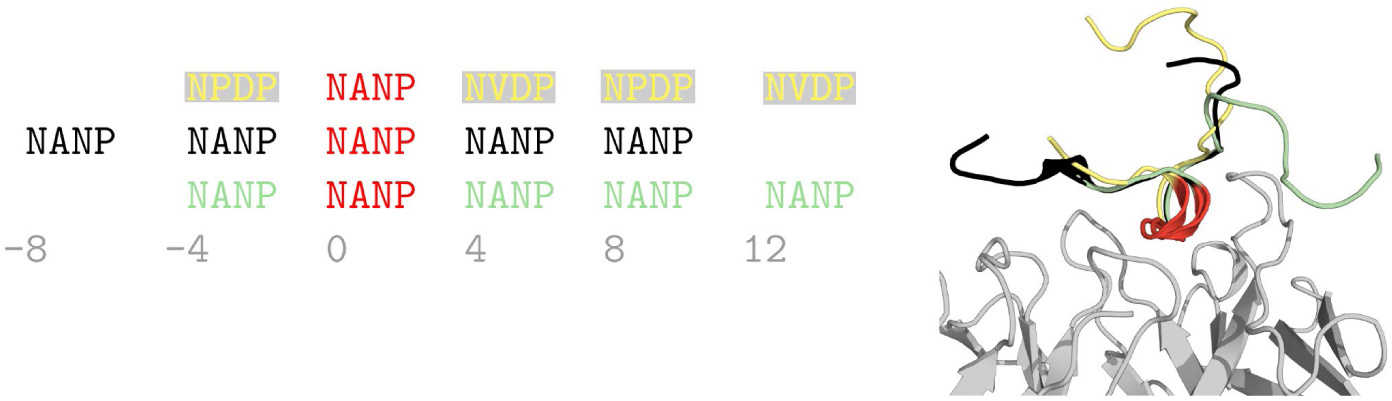
Positioning of epitope peptides in the antibody paratope. Left: the peptide sequences with residues centrally bound by the paratope coloured in red. Right: the corresponding cartoon representation of each sequence, with the peptides coloured according to the sequences and the antibody in grey. (Yellow: NPDP, Green: NANP with positioning like NPDP; Black: NANP modelled with one repeat added to both the N- and C-terminal sides).

## APPENDIX

### Supplementary Figures and Tables

**Figure S1.**
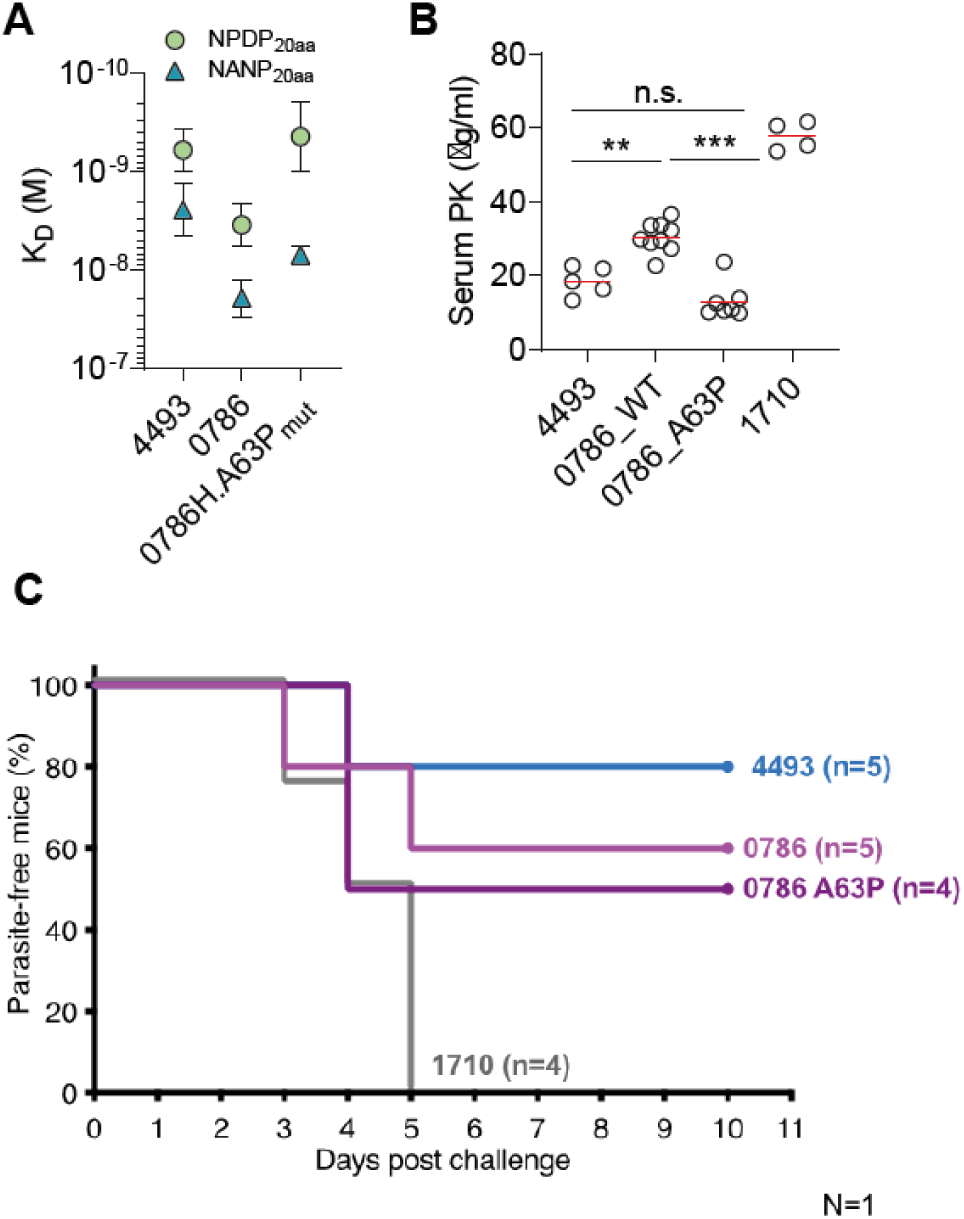
Affinity improving mutation H.A63P in mAb 0786 reduces mAb stability and does not enhance protective capacity. **A.** Affinities of mAb 0786 with and without H.A63P mutation compared to that of mAb 4493. **B.** Serum stability *in vivo* in mice at 20 h post intraperitoneal transfer of 150 µg of the indicated mAbs. **C.** Percentage parasite-free mice protected by a passive transfer of mAbs after a challenge with three bites of mosquitoes infected with transgenic PbPfCSP(mCherry) parasites. **P < 0.01, ***P < 0.001, n.s. not significant, two-tailed Mann-Whitney test. mAb 1710 was used as a negative control (Scally et al., 2018).

**Figure S2:**
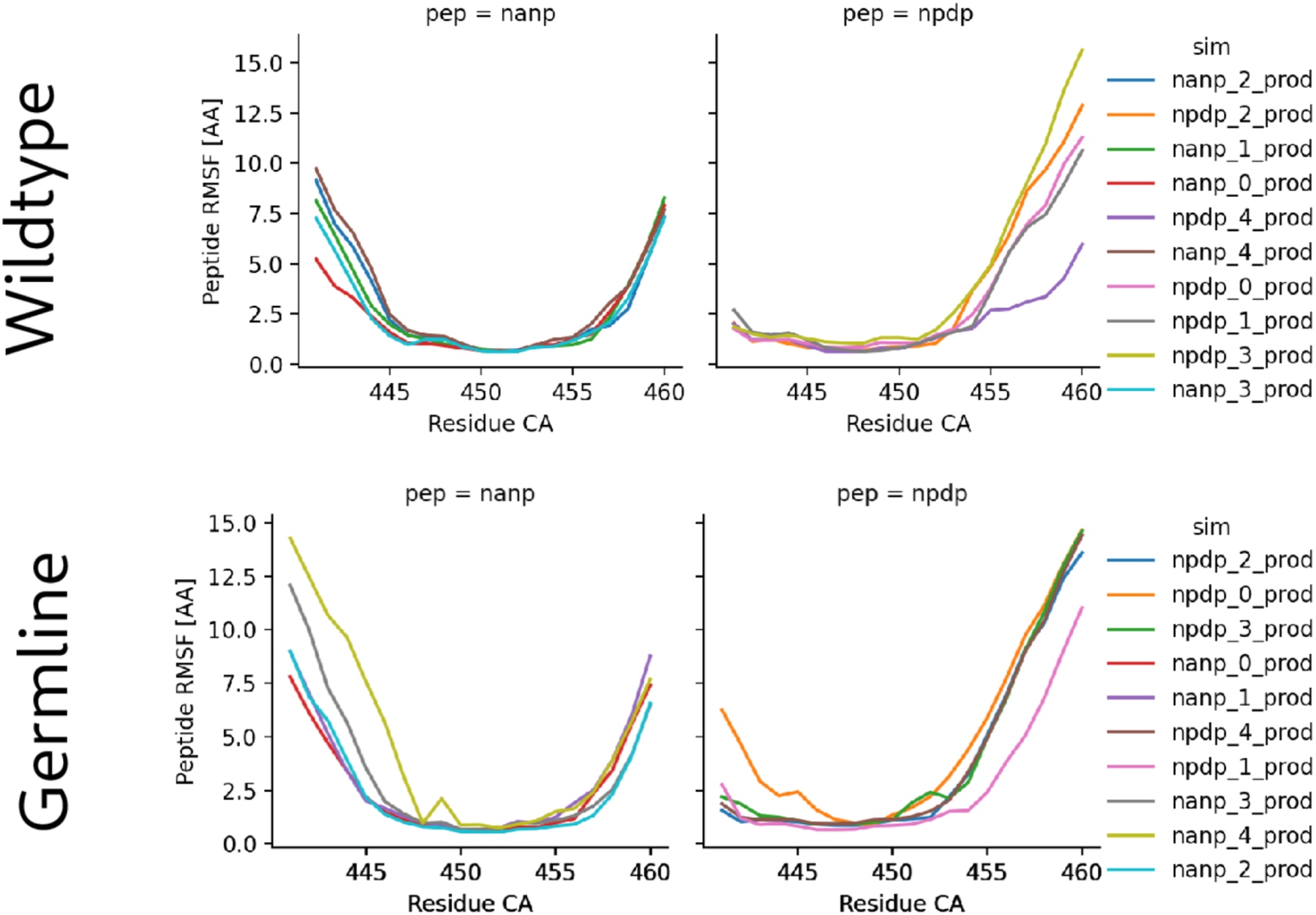
Fluctuations of the NANP and NPDP peptides bound to antibody during molecular dynamics simulations. The RMSF values were calculated for the Cα atoms of the peptide residues after alignment of the antibody residues of the trajectory frames onto their coordinates in the first frame. Note the decrease in N-terminal tail flexibility of the NANP peptide and larger standard deviation in RMSF between replicates for the NPDP C-terminal tail through maturation.

**Figure S3:**
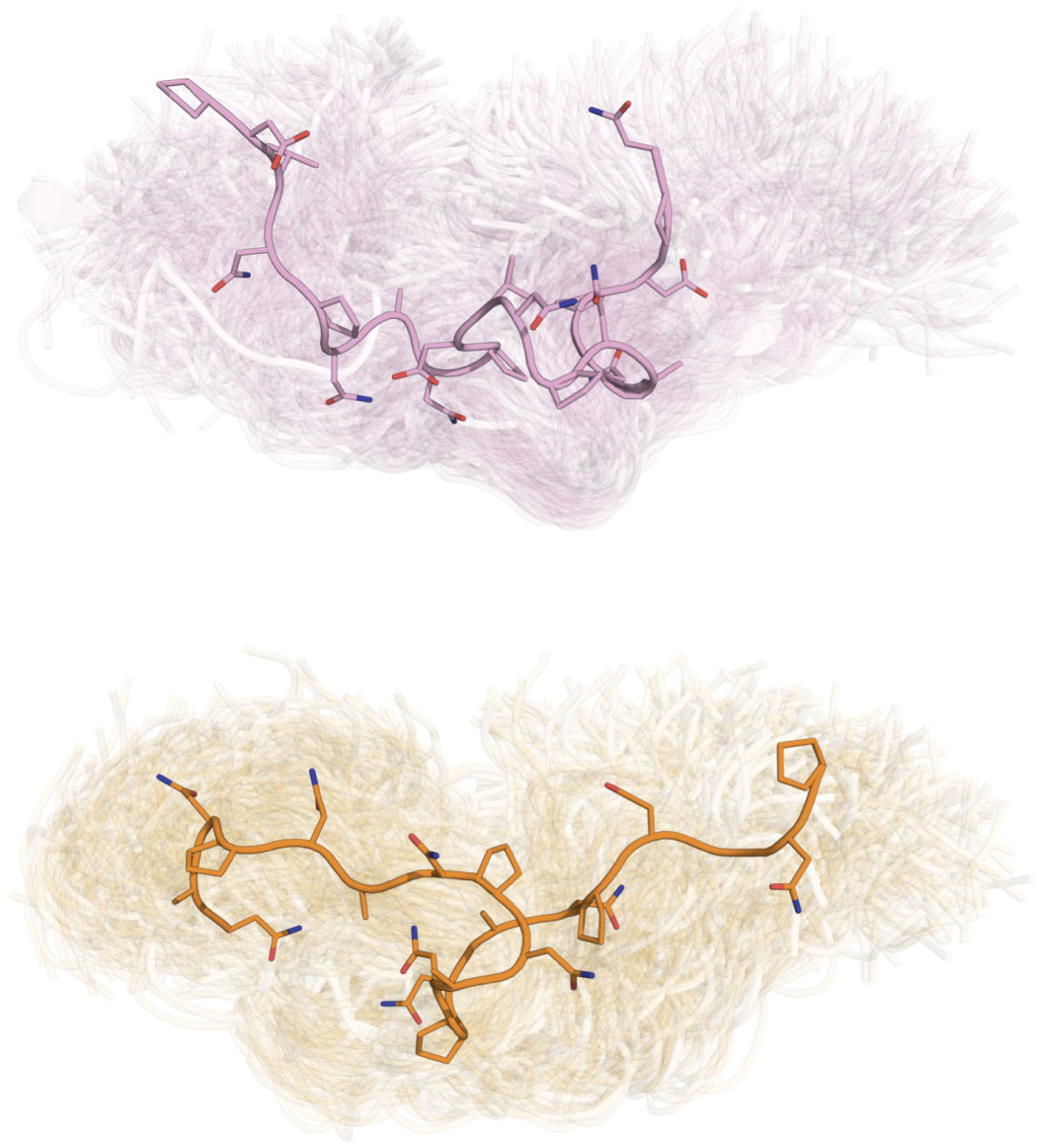
Unbound epitope peptides do not adopt a stable secondary structure. Conformations from conventional molecular dynamics (100ns, 10 frames/ns) simulations of the NPDP (pink) and NANP (orange) peptides in aqueous solution are shown superimposed with side chains in stick representation.

**Figure S4:**
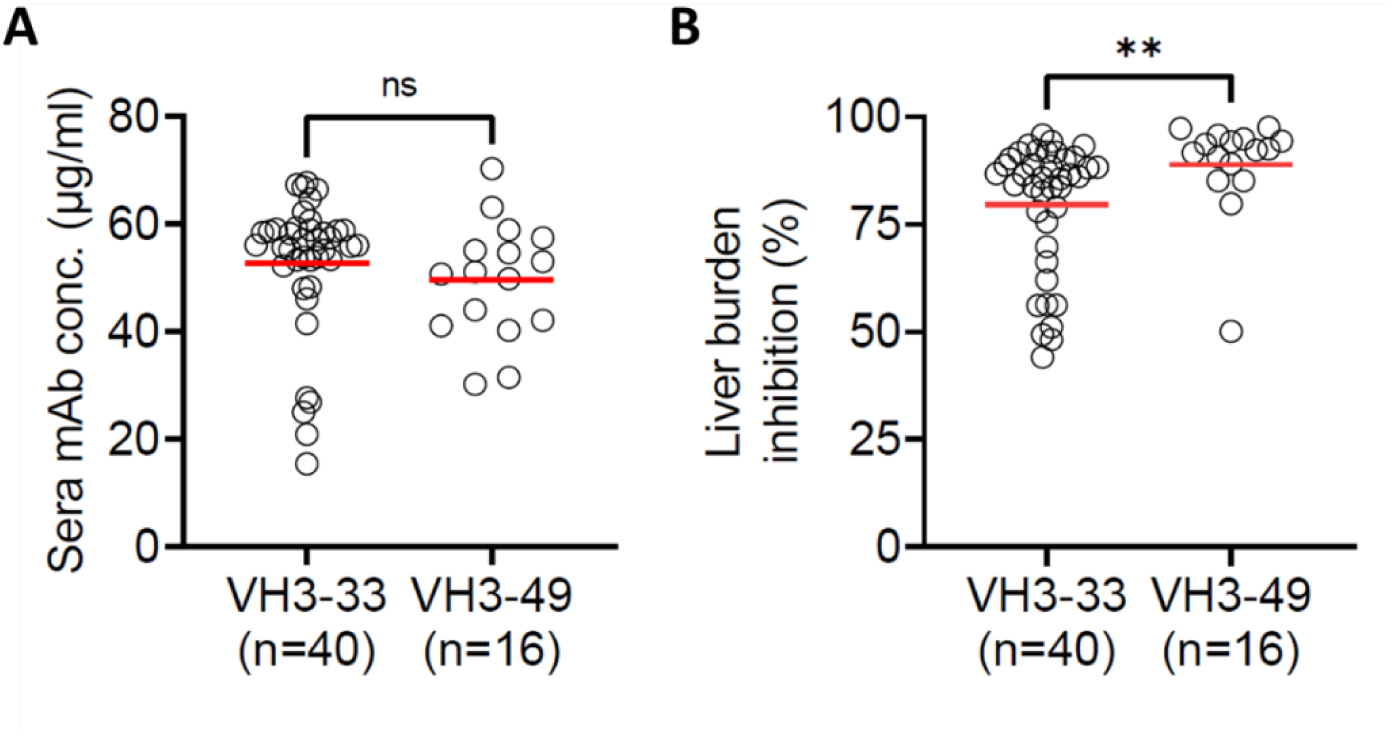
Comparison of VH3-33 and VH3-49 mAbs derived from RTS,S vaccination. *In vivo* serum stability in mice (A) and parasite inhibitory capacity measured as liver burden inhibition in mice (B) of VH3-33 and VH3-49 mAbs derived from RTS,S vaccinated individuals. Plots were made using the data published by Williams et al., 2024. Red lines indicate arithmetic mean. **P < 0.01, n.s. not significant, two-tailed Mann-Whitney test.

**Table S1:** Main settings of the equilibration protocol used for the conventional MD and RAMD simulations.

| | Equil. Step | t [ps] | Ensemble | Restraints [$\text{kJ mol}^{-1} \text{nm}^{-2}$] | Thermo/Barostat | $\tau$ [$\text{ps}^{-1}$] | T/p | T [K] | p [atm] |
| --- | --- | --- | --- | --- | --- | --- | --- | --- | --- |
| Conv. MD | 1 | 150 | NVT | Ab.: 4000; Pep.: 4000 | Nose-Hoover | 1.0 |  | 298.15 | – |
|  | 2 | 100 | NVT | Ab.: 2000; Pep.: 2000 | Berendsen | 1.0 |  | 298.15 | – |
|  | 3 | 100 | NPT | Ab.: 2000; Pep.: 2000 | Berendsen/Berendsen | 1.0/5.0 |  | 298.15 | 1.0 |
|  | 4 | 500 | NPT | Ab.: 1000; Pep.: 1000 | Nose-Hoover/Parrinello-Rahman | 1.0/5.0 |  | 298.15 | 1.0 |
|  | 5 | 500 | NPT | Ab.: 700; Pep.: 300 | Nose-Hoover/Parrinello-Rahman | 1.0/5.0 |  | 298.15 | 1.0 |
| τ-RAMD | 1 | 150 | NVT | Ab.: 4000 | Berendsen | 1.0 | | 298.15 | – |
|  | 2 | 100 | NPT | Ab.: 4000 | Berendsen/Berendsen | 1.0/5.0 |  | 298.15 | 1.0 |
|  | 3 | 50 | NPT | Ab.: 700 | Nose-Hoover/Parrinello-Rahman | 1.0/5.0 |  | 298.15 | 1.0 |
| | 4 | 100 ($\Delta t = 2$ fs) | NPT | – | Nose-Hoover/Parrinello-Rahman | 1.0/5.0 | | 298.15 | 1.0 |
| All Equil. Steps (unless indicated): $\Delta t = 1$ fs | | | | | | | | | |
| Energy Minimization steps: |  |  |  |  |  |  |  |  |  |
| | 1 | Steep, $E_{\text{tol}} = 1000 \text{ kJ mol}^{-1}$, max. 5000 steps, Restraints: Ab. $600 \text{ kJ mol}^{-1} \text{nm}^{-2}$ Pep. $40 \text{ kJ mol}^{-1} \text{nm}^{-2}$ | | | | | | | |
| | 2 | CG, $E_{\text{tol}} = 1000 \text{ kJ mol}^{-1}$, max. 5000 steps, Restraints: Ab. $40 \text{ kJ mol}^{-1} \text{nm}^{-2}$ Pep. $40 \text{ kJ mol}^{-1} \text{nm}^{-2}$ | | | | | | | |

Particle-Mesh Ewald and other settings were kept as in production simulations (see Methods).

**Table S2:** Conventional MD simulations generated for this study.

| System | Replica | $\sum$ Duration [ns] (10 frames/ns) |
| --- | --- | --- |
| Wildtype | 10 | 1000 |
| Germline | 5 | 500 |
| Wildtype w/o peptide | 5 | 500 |
| Wildtype capped peptide | 5 | 500 |
| Germline w/o peptide | 5 | 500 |
| Wildtype un-truncated | 5 | 500 |
| Germline un-truncated | 5 | 500 |
| H.N56Y<sub>rev</sub> | 5 | 500 |
| H.R59T<sub>rev</sub>-H.P63A<sub>rev</sub> | 5 | 500 |
| H.E102Y<sub>rev</sub>-H.S108A<sub>rev</sub> | 5 | 500 |
| NANP peptide | 2 | 162.9 |
| NPDP peptide | 2 | 200 |

Un-truncated simulations contained whole Fab fragments, while all other simulations contain only the variable regions.

### Positional ambiguity of NANP peptides

The positioning of the NPDP peptide is clear from the crystal structure due to the peptide’s non-palindromic four amino acid repeat sequence and therefore, the missing repeats can only be modelled on the C-terminal side (Murugan et al., 2020). On the other hand, from the crystal structure (PDB ID: 6o2c; (Murugan et al., 2020), the density map and the palindromic four aa repeat sequence, the positioning of the NANP repeat peptide in the paratope is ambiguous. The NANP peptide in complex with the mature wt antibody was therefore modelled with two distinct positions of the peptide to assess the impact of its positioning on the structure and dynamics. In one case, the NANP peptide was modelled with a positioning corresponding to that of the NPDP peptide (Fig. EV5, NANP: green; NPDP: yellow), and in the other case, the crystal structure of the NANP peptide was simply extended by one repeat on both the N- and C-terminal sides (Fig. EV5, black). For the GL_H+K_ and mutant antibodies, only models with the latter positioning of the peptide were generated.

### Uncharged epitope peptides

In the full-length PfCSP, both peptides simulated in this study would not be terminal and hence the terminal zwitterions present in the *in vitro* and *in silico* experiments at pH 7.5 constitute differences with respect to the full-length protein. For direct comparison between the *in vitro* and *in silico* experiments, these terminal charges were included in the computational models. However, to evaluate the possible influence of the terminal charges on antibody/peptide interactions, we built capped models without charged termini with PyMOL (The PyMOL Molecular Graphics System, Version 3.0 Schrödinger, LLC.) by prepending an acetyl group (ACE) to the N-terminus and appending an N-methyl (NME) to the C-terminus.

## Footnotes

1 https://github.com/HITS-MCM/gromacs-ramd

